# Increased efficacy of histone methyltransferase G9a inhibitors against MYCN-amplified Neuroblastoma

**DOI:** 10.1101/851105

**Authors:** Jacob Bellamy, Marianna Szemes, Zsombor Melegh, Anthony Dallosso, Madhu Kollareddy, Daniel Catchpoole, Karim Malik

**Author notes:** Joint first authors (Jacob Bellamy, Marianna Szemes).

## Abstract

Targeted inhibition of proteins modulating epigenetic changes is an increasingly important priority in cancer therapeutics, and many small molecule inhibitors are currently being developed. In the case of neuroblastoma (NB), a paediatric solid tumour with a paucity of intragenic mutations, epigenetic deregulation may be especially important. In this study we validate the histone methyltransferase G9a/EHMT2 as being associated with indicators of poor prognosis in NB. Immunological analysis of G9a protein shows it to be more highly expressed in NB cell-lines with *MYCN* amplification, which is a primary determinant of dismal outcome in NB patients. Furthermore, G9a protein in primary tumours is expressed at higher levels in poorly differentiated/undifferentiated NB, and correlates with high EZH2 expression, a known co-operative oncoprotein in NB. Our functional analyses demonstrate that siRNA-mediated G9a depletion inhibits cell growth in all NB cell lines, but, strikingly, only triggers apoptosis in NB cells with *MYCN* amplification, suggesting a synthetic lethal relationship between G9a and MYCN. This pattern of sensitivity is also evident when using small molecule inhibitors of G9a, UNC0638 and UNC0642. The increased efficacy of G9a inhibition in the presence of MYCN-overexpression is also demonstrated in the SHEP-21N isogenic model with tet-regulatable MYCN. Finally, using RNA sequencing, we identify several potential tumour suppressor genes that are reactivated by G9a inhibition in NB, including the *CLU, FLCN, AMHR2* and *AKR1C1-3*. Together, our study underlines the under-appreciated role of G9a in NB, especially in MYCN-amplified tumours.

## Introduction

Neuroblastoma (NB) is a biologically and clinically heterogeneous cancer arising from the developing sympathetic nervous system. About 25% of NB patients have a very poor prognosis clinical subtype characterized by amplification of the *MYCN* proto-oncogene (Brodeur et al. 1984; Brodeur 2003; Maris et al. 2007). Change of function gene mutations are relatively scarce in NB, but include the oncogene *ALK*, which is frequently mutated in familial NB and in up to 10% of sporadic cases (Mossë et al. 2008). This has prompted the notion that epigenetic aberrations are likely to contribute to NB pathogenesis.

Consistent with this hypothesis, evidence has accrued for the involvement of epigenetic modifiers, including histone methyltransferases (HMTs) in NB tumorigenesis. For example, we previously showed that knockdown of the HMTs EZH2, CARM1 or PRMT5 all decreased survival of NB cells (Park et al. 2015). High levels of the MYCN transcription factor leads to activation of survival/growth genes, but also repression of genes necessary for terminal differentiation in the sympathetic nervous system (Westermark et al. 2011; Gherardi et al. 2013). MYCN represses genes driving differentiation and apoptosis by a variety of means, including recruitment of epigenetic repressors, such as histone deacetylases (Iraci et al. 2011) and the Polycomb protein EZH2 (Corvetta et al. 2013). EZH2 has been independently shown to repress tumour suppressor genes in neuroblastoma, including *CASZ1, RUNX3, NGFR* and *CLU* (Wang et al. 2012). *CLU*, encoding Clusterin, has been characterized as a haploinsufficient tumour suppressor gene in NB (Chayka et al. 2009). The widespread involvement of HMTs in tumorigenesis has led to concerted pharmaceutical interest in developing selective inhibitors for HMTs (Helin and Dhanak 2013)

Another HMT implicated in NB is G9a (or EHMT2/KMT1C), the primary function of which is to mono- or di-methylate histone 3 lysine 9 (H3K9) (Tachibana et al. 2002; Shinkai and Tachibana 2011). These methylation marks are related to transcriptional activation and repression respectively, and G9a mediated gene silencing has been shown to be involved in regulating differentiation of embryonic stem cells (Wen et al. 2009). Moreover, G9a has multiple non-histone targets such as p53 (Huang et al. 2010) and chromatin remodelers Reptin and Pontin (Lee et al. 2010; Lee et al. 2011). In these cases, the post-translational methylation of proteins can either inactivate protein function, as in the case of p53, or in the case of Reptin and Pontin can direct these proteins to different targets to alter target gene expression. G9a can also act as a co-factor independent of its HMT activity by binding to nuclear receptor coactivator GRIP1 and forming a scaffold complex which in turn can activate downstream targets (Lee et al. 2006). G9a is known to be overexpressed in a variety of cancers such as colorectal (Zhang et al. 2014), bladder (Cho et al. 2011) and hepatocellular (Kondo et al. 2007) carcinomas suggesting it is an oncoprotein and therefore a viable therapeutic target for small molecule inhibitors. The G9a inhibitor BIX-01294 was one of the first HMT inhibitors discovered, displaying over 20-fold greater inhibition of G9a compared to the closely related HMT GLP (EHMT1). This inhibitor also showed no activity against a panel of HMTs including PRMT1, SUV39H1, SET7/9 and ESET, and was able to reduce demethylated H3K9 (H3K9me2) levels in chromatin (Kubicek et al. 2007). A second generation inhibitor UNC0638 is a potent and highly selective probe for G9a and GLP (>500-fold selectivity over other histone methyltransferases) and a high toxicity/function ratio of >100 (Vedadi et al. 2011). UNC0642 has improved pharmacokinetic properties and is suitable for use *in vivo* (Liu et al. 2013). UNC0638 and UNC0642 act as competitive substrate inhibitors, thus blocking the SET domain from acquiring methyl groups from its S-adenosyl-methionine (SAM) cofactor.

Three previous studies have alluded to the possibility of G9a as a therapeutic target in NB. On the basis of microarray database analysis, Lu et al proposed that G9a may be oncogenic in NB, and further showed that G9a knockdown or BIX-01294 treatment led to apoptosis in three NB cell-lines (Lu et al. 2013). In contrast, two other studies suggested that G9a knockdown or BIX-01294 treatment could trigger autophagic cell death (Ding et al. 2013; Ke et al. 2014, 2019), and that G9a-mediated epigenetic activation of serine-glycine metabolism genes is critical in oncogenesis. Taken together, these papers agree that inhibiting G9a may be beneficial for NB therapy, but the mode of action is unclear. In addition, the greatly more selective second generation of G9a inhibitors such as UNC0638 and UNC0642 have not been evaluated.

In this study, we comprehensively assess the association of G9a with key prognostic factors in NB, specifically differentiation status and MYCN over-expression. We further evaluate UNC0638 and UNC0642 as potential therapeutic agents for NB, and identify putative tumour suppressor genes that are repressed by G9a in NB. Our data strongly suggest that G9a inhibition may be especially beneficial for poor-prognosis NB driven by *MYCN* amplification.

## Materials and Methods

### Neuroblastoma cell lines and culture conditions

Neuroblastoma cell lines were kindly supplied by Prof. Deborah Tweddle (Newcastle University), Prof. Manfred Schwab (German Cancer Research Center), and Robert Ross (Fordham University), the Childrens Oncology Group (Texas Tech University Health Sciences Center) or purchased from Deutsche Sammlung von Mikroorganismen und Zellkulturen (DSMZ). Cell lines were cultured in Dulbecco’s modified eagle’s medium (DMEM):F12□HAM (Sigma) supplemented with 10% (v/v) foetal bovine serum (FBS) (Life technologies), 2mM L-glutamine, 100 U/mL penicillin, 0.1 mg/mL streptomycin, and 1% (v/v) non-essential amino acids. SH-EP□Tet21N cells were cultured in RPMI 1640 (Gibco), supplemented with 10% (v/v) tetracycline-free FBS (Life technologies), 2 mM L-Glutamine, 100 U/mL penicillin, 0.1 mg/mL streptomycin, and 1μg/mL tetracycline. Cell counts and cell viability were assessed using Countess automated cell counter and trypan blue (Thermo Fisher Scientific). Transient knockdowns were performed by using short interfering RNA (siRNA), targeting *G9a/EHMT2* (5’-GAACAUCGAUCGCAACAUCdTdT-3’/5’-GAUGUUGCGAUCGAUGUUCdTdT-3’) in a reverse transfection protocol, with 50 nM siRNA and Lipofectamine RNAiMAX (Invitrogen), both diluted in OptiMEM media (Invitrogen). Non-targeting siRNAs were used as control (5’-UGGUUUACAUGUUUUCUGAdTdT-3’/5’-UCAGAAAACAUGUAAACCAdTdT-3’). For G9a inhibition, attached cells were treated with BIX-01294 (Tocris), UNC0638, (Tocris) and UNC0642 (Tocris) dissolved in DMSO, at the indicated concentrations.

### MTT cell viability assay

NB cells were seeded in 96 well plates and treated the next day in triplicate with a serial dilution of UNC0638/0642. After 72 hours, we added 10μL of MTT (5 mg/mL) (Sigma), followed by 50μL of SDS lysis buffer (10% SDS (w/v), 1/2500 (v/v) 37% HCl) after a further 3 hours. Following an overnight incubation at 37°C, the plates were read at 570 and 650 nm, using SpectraMax 190 plate reader (Molecular Devices).

### Protein Extraction and Western blot

Floating and attached cells were lysed in Radioimmunoprecipitation assay (RIPA) buffer. Protein concentration was determined by using Micro BCA TM protein assay kit (Thermo Fisher). Immunoblotting was performed as described previously (Park et al. 2015). The following antibodies were used to detect G9a (ab185050, Abcam), cPARP (ab32064, Abcam), MYCN (B8.48, Santa Cruz, SC-53993), cCaspase 3 (9664, Cell Signaling Technology), LC3B (L7543, Sigma), and β-Actin (A3854, Sigma), according to manufacturer’s instructions.

### RNA extraction, reverse transcription and qPCR

RNA was extracted from attached cells by using RNeasy Plus or miRNeasy kits (QIAGEN) according to manufacturer’s instructions and subsequently transcribed into cDNA with Superscript IV (Invitrogen). Quantitative PCR was performed by using QuantiNova kit on Mx3500P PCR machine (Stratagene). The following oligonucleotide primers were used to detect target gene expression: *AMHR2* F-TACTCAACCACAAGGCCCAG, R-GGTCTGCATCCCAACAGTCT, *FLCN* F-TCTCTCAGGCTGTGGGAGC R-CCAGCATGCGGAAAGAAG, *AKR1C1* F-CCTAAAAGTAAAGCTTTAGAGGCCACC, R-GAAAATGAATAAGGTAGAGGTCAACATAAT, *AKR1C2*, F-CCTAAAAGTAAAGCTCTAGAGGCCGT, R-GAAAATGAATAAGATAGAGGTCAACATAG, *AKR1C3* F-CTGATTGCCCTGCGCTAC, R-TCCTCTGCAGTCAACTGGAAC, *CLU* F-AGCAGCTGAACGAGCAGTTT, R-AGCTTCACGACCACCTCAGT, *TBP* F-AGCCACGAACCACGGCACTGAT, R-TACATGAGAGCCATTACGTCGT, *ALK* F-CGACCATCATGACCGACTACAA, R-CCCGAATGAGGGTGATGTTTT.

### Cell cycle analysis

Propidium-iodide labelling and fluorescence activated cell sorting analysis to detect cell cycle phases was performed as previously described (Park et al. 2015).

### Immunohistochemistry

Tissue microarrays (TMAs), containing 50 peripheral neuroblastic tumours were stained using antibodies for EZH2 (NCL-L-EZH2, Novocastra) and G9a/EHMT2 (EPR4019(2), Abcam). Immunohistochemistry was independently scored by two pathologists blinded to the specimens, and a score of 1-4 was assigned based on proportion of positive cells (0, no staining; 1, sporadic staining of individual cells; 2, <20% of cells stained; 3, 20% to 50% of cells stained; 4, >50% of cells stained). All human tissues were acquired with appropriate local research ethics committee approval. Immunohistochemistry was performed with a Leica Microsystem Bond III automated machine using the Bond Polymer Refine Detection Kit (Ref DS9800) followed by Bond Dab Enhancer (AR9432). The slides were dewaxed with Bond Dewax Solution (AR9222). Heat mediated antigen retrieval was performed using Bond Epitope Retrieval Solution for 20 mins.

### RNA-seq and bioinformatic analysis

LAN-1 cells were treated with 3µM BIX-01294 and DMSO vehicle as control for 72 hours and were subsequently harvested. RNA was extracted by using miRNeasy Mini Kit (Qiagen), according to manufacturer’s instructions. Libraries were constructed and sequenced as previously described (Szemes et al. 2018). Briefly, cDNA libraries were prepared from 1 ug RNA (TruSeq Stranded Total RNA Library Prep Kit, Illumina) and 100 bp, paired end reads were sequenced on Illumina HiSeq 2000. The reads were aligned to the human genome (hg38) by using TopHat2 (v2.0.14)and the alignment files (BAM) files were further analysed in SeqMonk v1.45. (https://www.bioinformatics.babraham.ac.uk/projects/seqmonk/). Gene expression was quantified by using the Seqmonk RNA-seq analysis pipeline. Differentially expressed genes (DEG) were identified by DESEQ2 (p<0.005). sets, and a minimum fold difference threshold of 1.3 was applied. RNA sequencing data is available from the European Nucleotide Archive (ENA) under the study accession number PRJEB35417. We performed Gene Signature Enrichment Analysis (GSEA) on a preranked list of log2-transformed relative gene expression values (Broad Institute). Kaplan Meier survival analysis, indicating the prognostic value of the expression of genes or metagenes was performed by using the Kaplan scan tool in R2 Genomics Analysis and Visualization Platform (http://r2.amc.nl).

## Results

### G9a expression correlates with poor prognosis and MYCN amplification in NB

We first assessed the expression of *G9a/EHMT2* in an RNA-seq dataset of 498 primary NBs (the SEQC dataset, GSE62564) (Su et al. 2014) using the R2: Genomics Analysis and Visualization Platform (http://r2.amc.nl). Kaplan-Meier analysis of overall survival showed that high *G9a*/*EHMT2* expression was significantly associated with poor survival, (Figure 1A). Moreover, *G9a/EHMT2* has a significantly increased expression in MYCN-amplified (MNA) NBs (Figure 1B). In order to assess whether this relationship at the RNA level was also apparent at the protein level, we conducted immunoblotting of G9a protein expression in NB cell lines with and without MNA, and confirmed that increased G9a protein expression was apparent in MNA NB cell lines (Figure 1C-D). We next examined G9a protein expression in a tissue microarray of 50 primary neuroblastic tumours, in parallel with EZH2, which is known to be involved in NB. Strikingly, expression of G9a and EZH2 was barely detectable in the more differentiated ganglioneuromas and ganglioneuroblastomas, generally regarded as low risk tumours (Figure 2A-B). However, high nuclear expression of both G9a and EZH2 was observed in the poorly differentiated and undifferentiated tumours, which are commonly associated with higher risk and poor outcome (Figure 2 D). Accordingly, G9a immunopositivity strongly correlated with the differentiation status of neuroblastic tumours (Figure 2E). The few G9a positive cells in the more differentiated tumours were all undifferentiated neuroblasts, underlining the link between differentiation status and G9a expression. Scoring the tumours for G9a and EZH2 expression according to the percentage of immunopositive cells revealed that G9a and EZH2 scores were very highly correlated (R=0.76, p=1.45e-10, Figure 2F), indicative of a potential functional interplay between these HMTs. Unfortunately, MYCN status was available for only a few tissue microarray samples, therefore not sufficient for statistical analysis. Nevertheless, our expression analyses at the RNA and protein levels shows that G9a over-expression is associated with poor prognosis NB and MYCN/EZH2 status.

**Figure 1.**
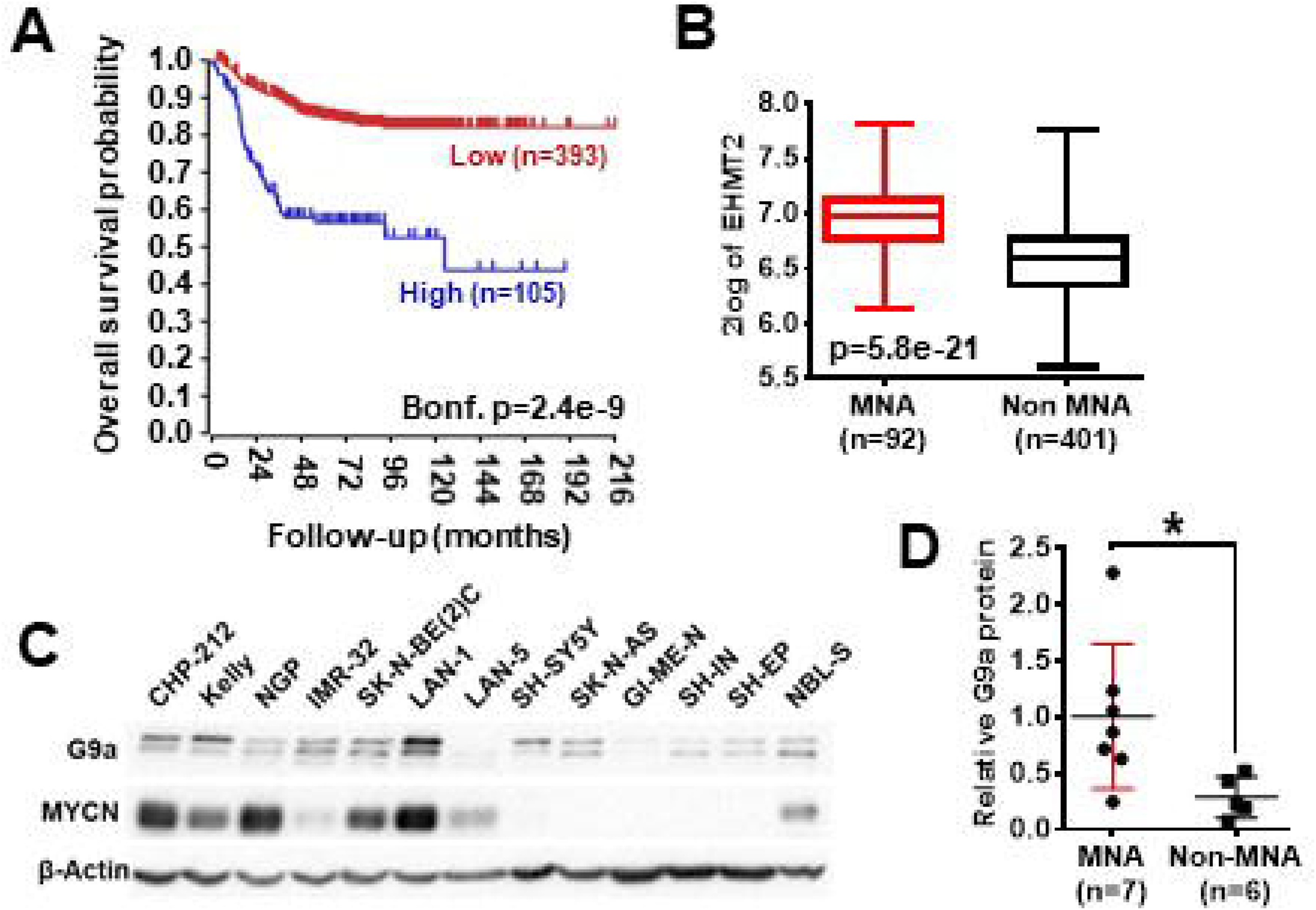
G9a mRNA and protein expression correlate with poor prognosis and MYCN amplification. **(A)** Kaplan Meier analysis showing that high expression of G9a correlates with poor prognosis in primary NB (SEQC, GSE62564). Bonferroni-corrected p values of log rank test are shown. **(B)** Significantly higher G9a mRNA expression is observed in MNA neuroblastoma relative to non MNA. The p-values are calculated by one-way ANOVA. **(C)** Immunoblot of G9a protein expression in a panel of MYCN-amplified and non-amplified NB cell lines. β-Actin is used as a loading control. Representative of n=3. **(D)** Scatter dot plot of relative G9a protein is generated from semi-quantitative densitometry of blots from **(C)**. G9a protein expression is higher in MNA neuroblastoma when normalised to β-Actin. Significance measured by unpaired T test (* p<0.05).

**Figure 2.**
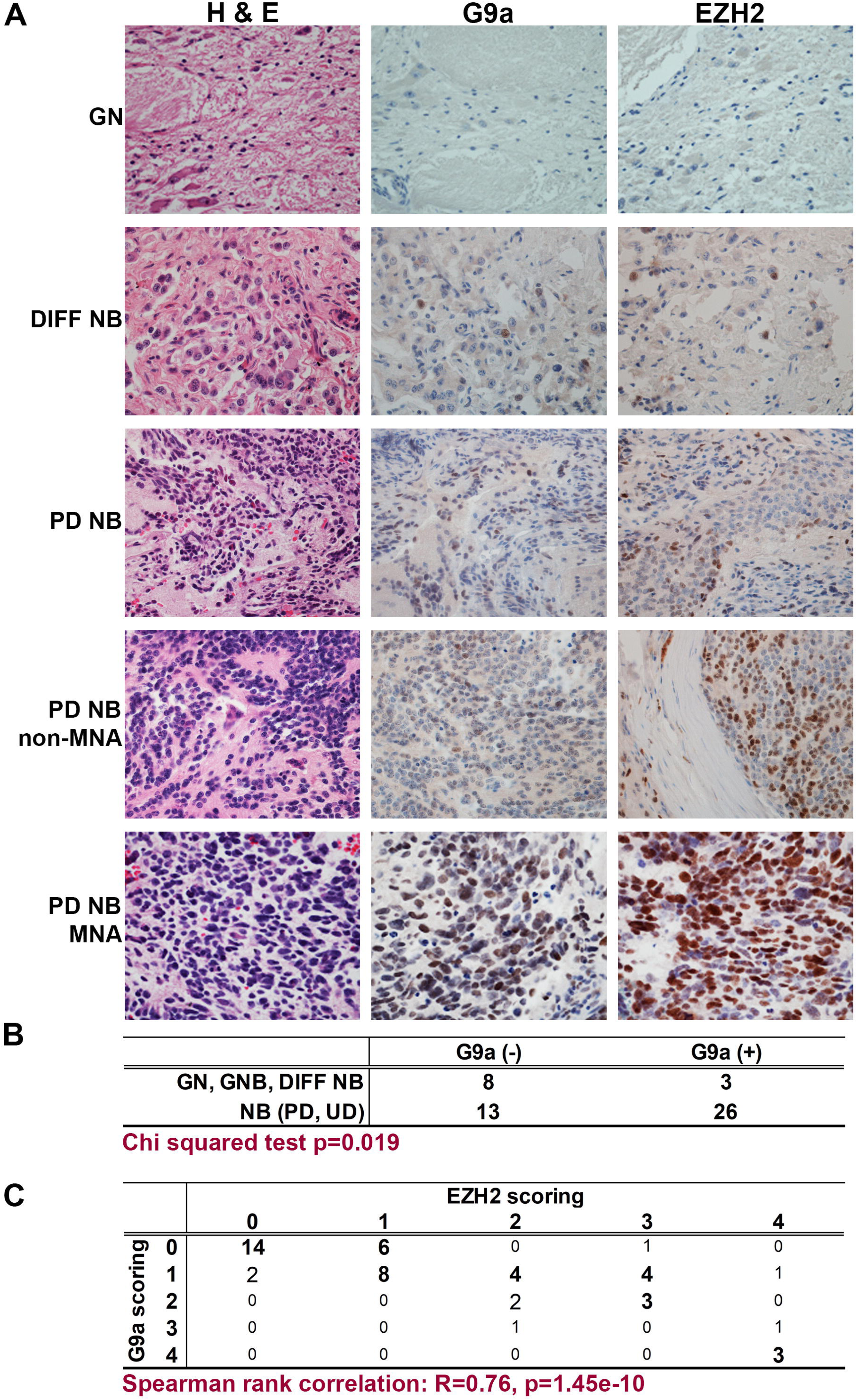
Immunohistochemical detection of G9a and EZH2 in primary NBs. **(A)** Haematoxylin and Eosin staining, G9a and EZH2 immunohistochemistry of neuroblastic tumours. (GN=ganglioneuroma, DIFF NB= differentiating neuroblastoma, PD NB=poorly differentiated NB, UD NB=undifferentiated NB, MNA=MYCN amplified). **(B)** Positive staining for G9a correlates with INPC differentiation status of neuroblastic tumours. **(C)** Immunopositivity was scored based on the proportion of positive cells (0, no staining; 1, sporadic staining of individual cells; 2, <20% of cells stained; 3, 20% to 80% of cells stained; 4, >80% of cells stained). The proportion of G9a and EZH2 positive cells strongly and significantly correlated in neuroblastic tumours.

### Short interfering RNA mediated G9a depletion leads to apoptotic cell death in MNA neuroblastoma cells

As there is increased expression of G9a in MNA neuroblastoma cell lines, we next evaluated the effect of short□interfering RNA (siRNA) mediated G9a depletion on three MNA cell lines and three non-MNA cell lines. Quantification of adherent and floating cells following knockdown of G9a in MNA Kelly, LAN-1, and SK-N-BE(2)C showed a consistent increase in the percentage of dead cells (p<0.05) and decrease in the number of live cells in the population (Figure 3A). In order to assess the mode of reduced cell survival, immunoblotting was carried out with markers for apoptosis (cleaved PARP and cleaved caspase 3) and autophagy (LC3B). In all three MNA lines, apoptotic cell death was verified by the increase in apoptosis markers (Figure 3A, lower panel). In contrast, no increases in LC3B were apparent, suggesting little or no effect on autophagy. Knockdown in the presence of QVD, a caspase inhibitor, led to decreased floating cells after G9a knockdown, and, as expected, no apoptotic markers (Supplementary Figure 1A-B). MYCN protein also decreased considerably after G9a knockdown in all three MNA lines. Cell cycle analysis of Kelly cells after G9a knockdown demonstrated a significant increase of cells in G1 and a decrease of cells entering S-phase (Supplementary Figure 1C).

**Figure 3.**
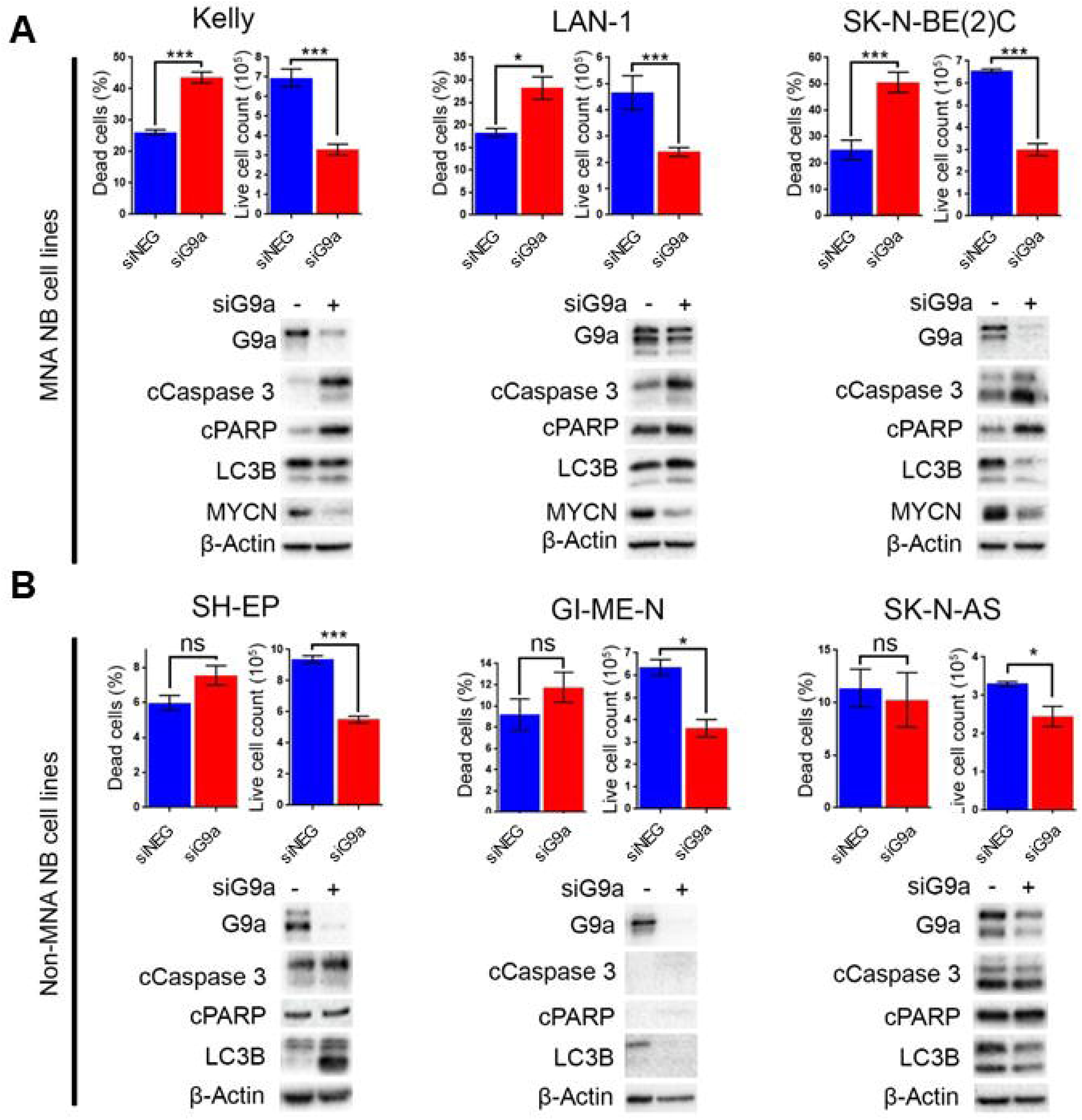
Apoptotic cell death following G9a depletion is dependent on MYCN. **(A)** Floating and adherent cells from MNA cell lines were harvested and counted by trypan blue inclusion assay following G9a depletion. The left-hand graph of each cell line shows the percentage dead cells between the G9a depleted and negative control. The right-hand graph shows the live cell count of the cells harvested, which is used as a proxy for cell growth. Significant changes are measured the asterisks (* p<0.05, *** p<0.01, ns not significant, n=3). Error bars show the SEM. For each cell line, an immunoblot showing effective depletions, apoptosis and autophagy markers is shown below. β-Actin is used as loading control. The blots are representative of n=3. **(B)** Live and dead cell counts and Western blots in non-MNA cell lines.

In contrast to these effects in MNA cell-lines, G9a depletion-associated cell death was not seen in the three non-MNA cell lines tested (SH-EP, GI-ME-N, and SK-N-AS), as there was no significant increase in the percentage of dead cells seen following the depletion. Despite this, decreased proliferation of these lines was indicated by a significant decrease in the number of live cells (Figure 3B). The observed lack of cell death was confirmed by immunoblots showing no change in apoptotic markers cleaved PARP and caspase 3 (Figure 3B). Variable effects on LC3B were observed, increasing after knockdown in SH-EP cells and decreasing in GI-ME-N and SK-N-AS cell-lines. Taken together, our analysis of NB cell lines strongly suggests a requirement on G9a for cell survival of MNA NB.

To confirm the G9a dependency of MYCN over-expressing NB cells, G9a depletion was evaluated in isogenic SH-EP□Tet21N (S21N) cells with Tet-inducible MYCN expression. With MYCN induced, G9a depletion led to a significant increase in the percentage of dead cells, whereas there was no change in cells without MYCN induction (Figure 4A). As before, this was despite a significant decrease in live cell count that occurred across both induced and non-induced MYCN S21N cells (Figure 4A). This was further confirmed by the immunoblots showing that G9a depletion led to an increase in apoptosis markers in the MYCN induced cells only (Figure 4B). These experiments further emphasise the G9a dependency of MYCN-overexpressing NB cells.

**Figure 4.**
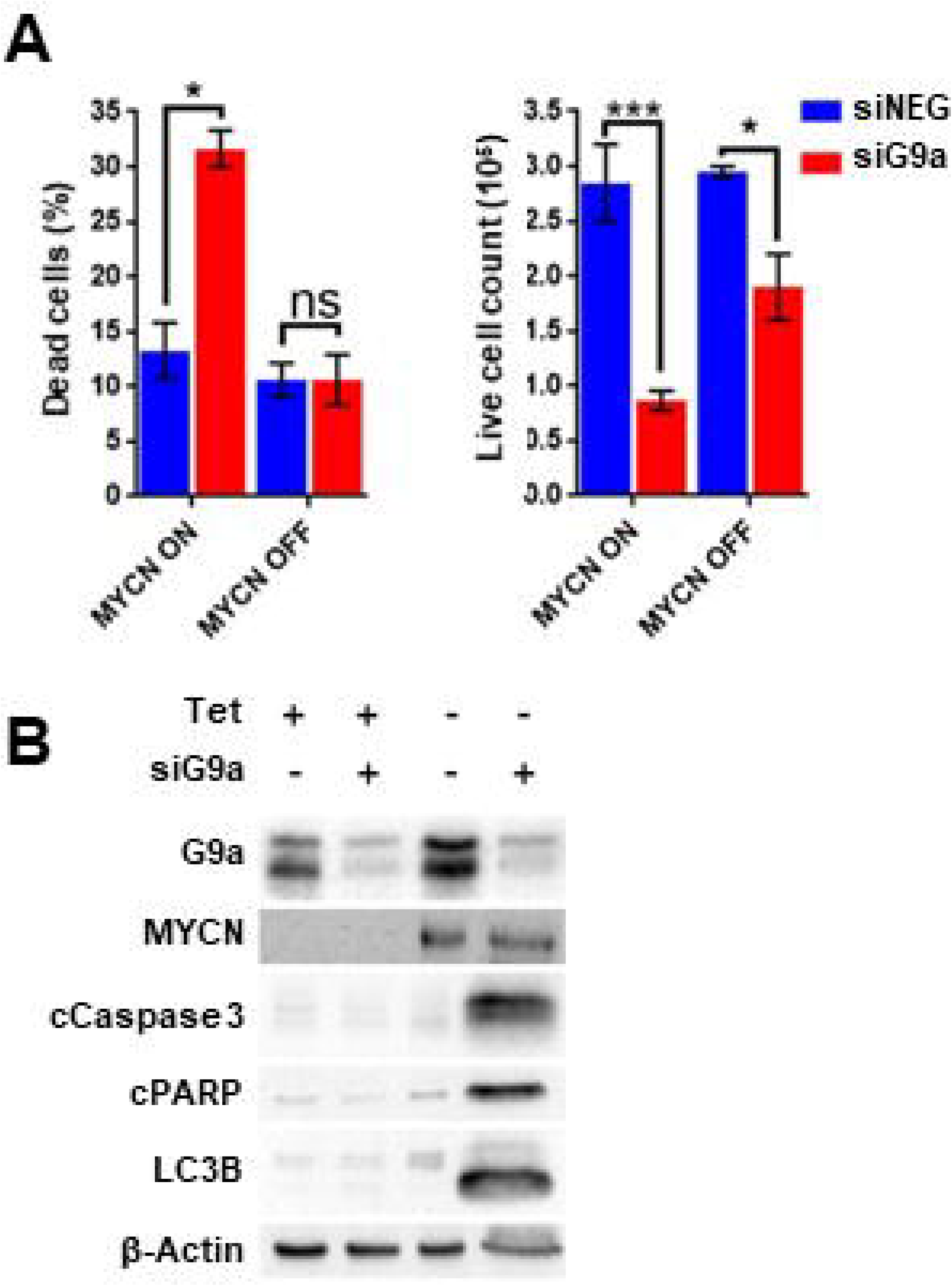
G9a depletion leads to cell death in MYCN induced S21N cells only. **(A)** Floating and adherent cells from S21N cells, with and without MYCN induction, were harvested and counted by trypan blue inclusion assay following G9a depletion as previously. **(B)** Immunoblot of G9a and apoptosis and autophagy markers.

### MNA NB cell are more sensitive to G9a inhibitors UNC0638 and UNC0642

As our genetic interference analyses demonstrate that NB cells may be vulnerable to decreased G9a activity, we treated a panel of 13 NB cell lines and 2 two disease-free control cell lines with G9a SMIs and conducted cell survival assays. Whilst all NB cell lines exhibited reduced viability upon treatment with UNC0638 (Figure 5A) and UNC0642 (Figure 5B) in a concentration dependent manner, there was no effect on the disease-free lines RPE-1 and NF-TERT. Importantly, consistent with our genetic interference data, a clear pattern of greater sensitivity of MNA lines was observed. For UNC0638, the average IC_50_ was 8.3μM for MNA lines, compared to 19μM for non-MNA lines (p<0.01). Similarly, for UNC0642 the average IC_50_s were 15μM and 32μM for MNA and non-MNA lines respectively (Figure 5C-F).

**Figure 5.**
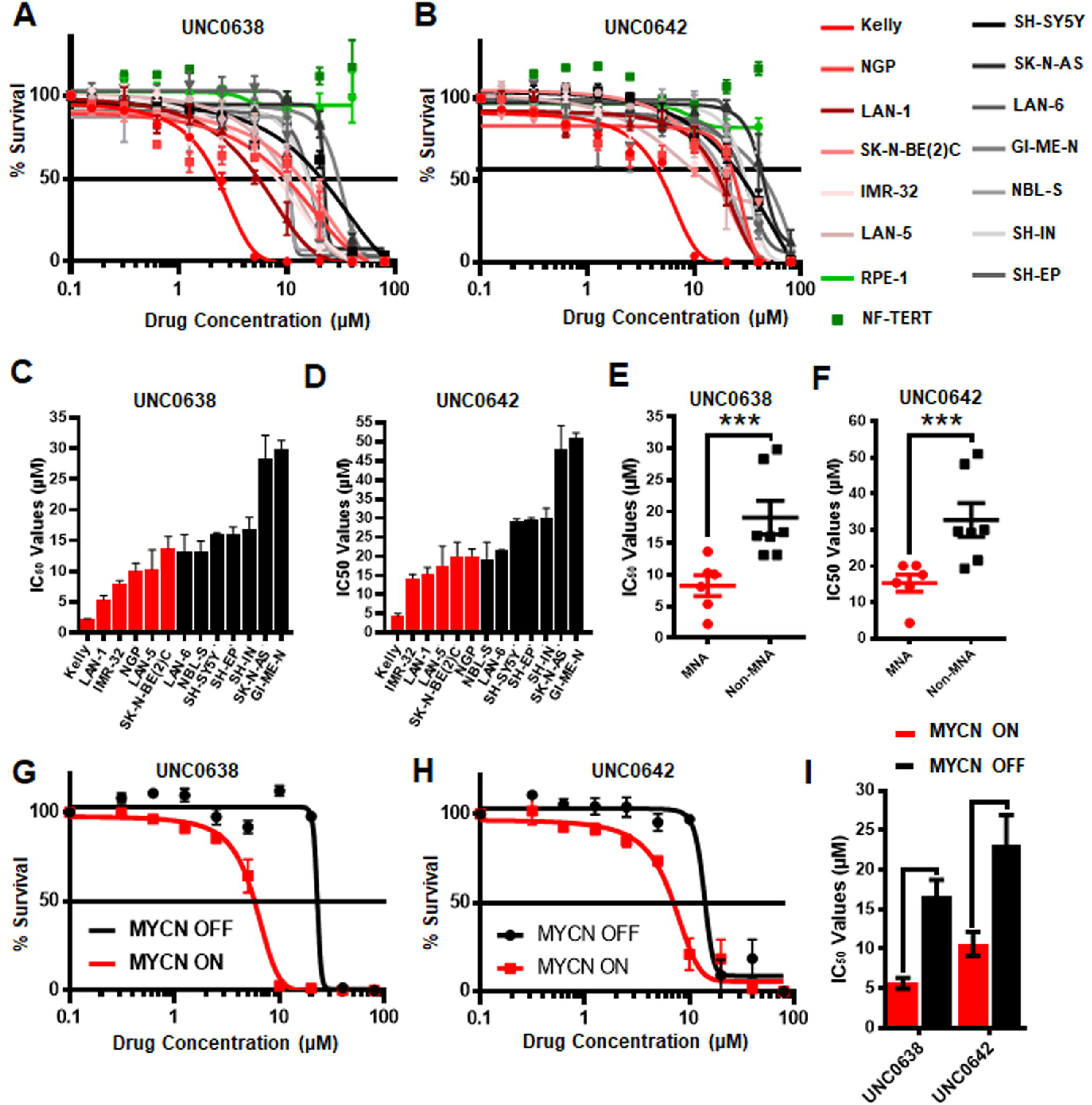
Sensitivity of neuroblastoma cell-lines to G9a inhibitors UNC0638 and UNC0642. **(A)** Thirteen neuroblastoma cell lines, including six MYCN-amplified (shades of red lines), seven non-MYCN-amplified (shades of black lines), and two non-cancerous cell lines (shades of green lines) were screened by MTT based cell proliferation assay after 72 hours to determine sensitivity to G9a SMI UNC0638. 0.0 μM was plotted as 0.1 μM, to be able to chart on log scale. Error bars show SEM. n≥ 3. **(B)** MTT assay of 13 NB and two normal cell lines using G9a SMI UNC0642. **(C)** Bar chart of IC_50_ values for UNC0638 with all cell lines. Error bars show SEM. **(D)** Bar chart of IC_50_ values with UNC0642. Error bars show SEM. **(E)** MTT assay based IC_50_ values for UNC0638 from **(A)** are visualized as a scatterplot between MNA and non-MNA cell lines. Error bars show the SEM. *** p<0.01, unpaired t test. **(F)** - MTT assay based IC_50_ values for UNC0642 from **(B)** are visualized as a scatterplot between MNA and non-MNA cell lines. Error bars show the SEM. *** p<0.01, unpaired t test. **(G)** S21N cell line with and without induced MYCN were screened by MTT based cell proliferation assay after 72 hours to determine sensitivity to G9a inhibitors UNC0638. 0.0 μM was plotted as 0.1μM to chart on log scale. Error bars show SEM. N=3. **(H)** MTT assay of induced and uninduced S21N cells using UNC0642. **(I)** – Bar chart of S21N IC_50_ values with and without induction. Error bars show the SEM. *** p<0.01, unpaired t test.

Next, UNC0638 and UNC0642 were evaluated in S21N cells to further assess the association of MYCN over-expression and sensitivity to pharmaceutical G9a inhibition. S21N cells with induced (high) MYCN were more sensitive to G9ai by UNC0638 and UNC0642 than the uninduced cells (Figure 5G-H). As with the cell-line panel, UNC0638 was slightly more potent than UNC0642, with the IC_50_ values for MYCN induced cells being 5.7μM and 10.6μM respectively, compared to 16.6μM and 23.2μM for uninduced S21N cells (Figure 5I). All the IC_50_ values for the cell-lines analysed are detailed in Table 1. Thus, our pharmaceutical inhibition assays reflect our genetic interference data, showing that NB cell-lines are sensitive to both UNC0638 and UNC0642, with MYCN over-expressing cells being significantly more sensitive than non-MNA lines.

**Table 1.**
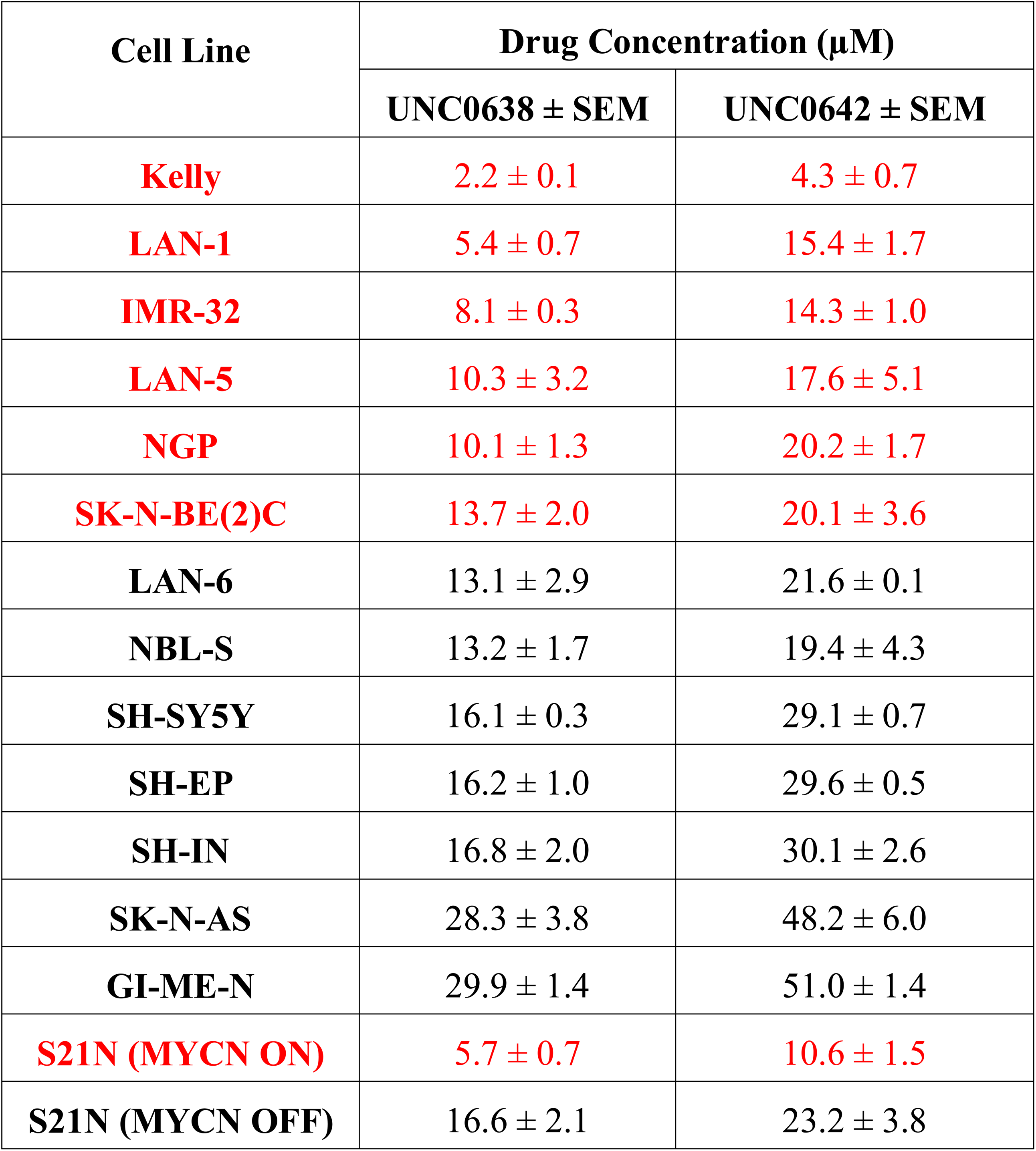
IC_50_ values of NB cell lines with G9a inhibitors UNC0638 and UNC0642. Values are means ± SEM of n≥ 3. MNA cell lines are coloured red.

| Cell Line | Drug Concentration ($\mu\text{M}$) | |
| --- | --- | --- |
| | UNC0638 ± SEM | UNC0642 ± SEM |
| **Kelly** | 2.2 ± 0.1 | 4.3 ± 0.7 |
| **LAN-1** | 5.4 ± 0.7 | 15.4 ± 1.7 |
| **IMR-32** | 8.1 ± 0.3 | 14.3 ± 1.0 |
| **LAN-5** | 10.3 ± 3.2 | 17.6 ± 5.1 |
| **NGP** | 10.1 ± 1.3 | 20.2 ± 1.7 |
| **SK-N-BE(2)C** | 13.7 ± 2.0 | 20.1 ± 3.6 |
| **LAN-6** | 13.1 ± 2.9 | 21.6 ± 0.1 |
| **NBL-S** | 13.2 ± 1.7 | 19.4 ± 4.3 |
| **SH-SY5Y** | 16.1 ± 0.3 | 29.1 ± 0.7 |
| **SH-EP** | 16.2 ± 1.0 | 29.6 ± 0.5 |
| **SH-IN** | 16.8 ± 2.0 | 30.1 ± 2.6 |
| **SK-N-AS** | 28.3 ± 3.8 | 48.2 ± 6.0 |
| **GI-ME-N** | 29.9 ± 1.4 | 51.0 ± 1.4 |
| **S21N (MYCN ON)** | 5.7 ± 0.7 | 10.6 ± 1.5 |
| **S21N (MYCN OFF)** | 16.6 ± 2.1 | 23.2 ± 3.8 |

### UNC0638 leads to increased apoptosis specifically in MNA NB lines

G9a knockdown analyses demonstrated that only MNA NB lines underwent programmed cell death, despite all lines showing some degree of growth inhibition. To determine if the effects on cell death and proliferation following G9a drug inhibition are akin to the results following G9a depletion as shown in Figure 3, three MNA and three non-MNA NB cell lines were treated with UNC0638 and cell growth effects quantified, together with an assessment of apoptosis and autophagy markers. The three most sensitive three MNA lines, Kelly, IMR-32 and LAN-1 all showed a significant increase in the percentage of dead cells following treatment (Figure 6A). Apoptosis was verified by immunoblotting, all 3 treated lines showing an increase in cleaved PARP and caspase 3 (Figure 6A, lower panel). Although Kelly and LAN-1 cells showed some increase in LC3B, IMR-32 cells exhibited a decrease of LC3B. As with knockdowns, all 3 MNA lines showed a decrease in MYCN protein.

**Figure 6.**
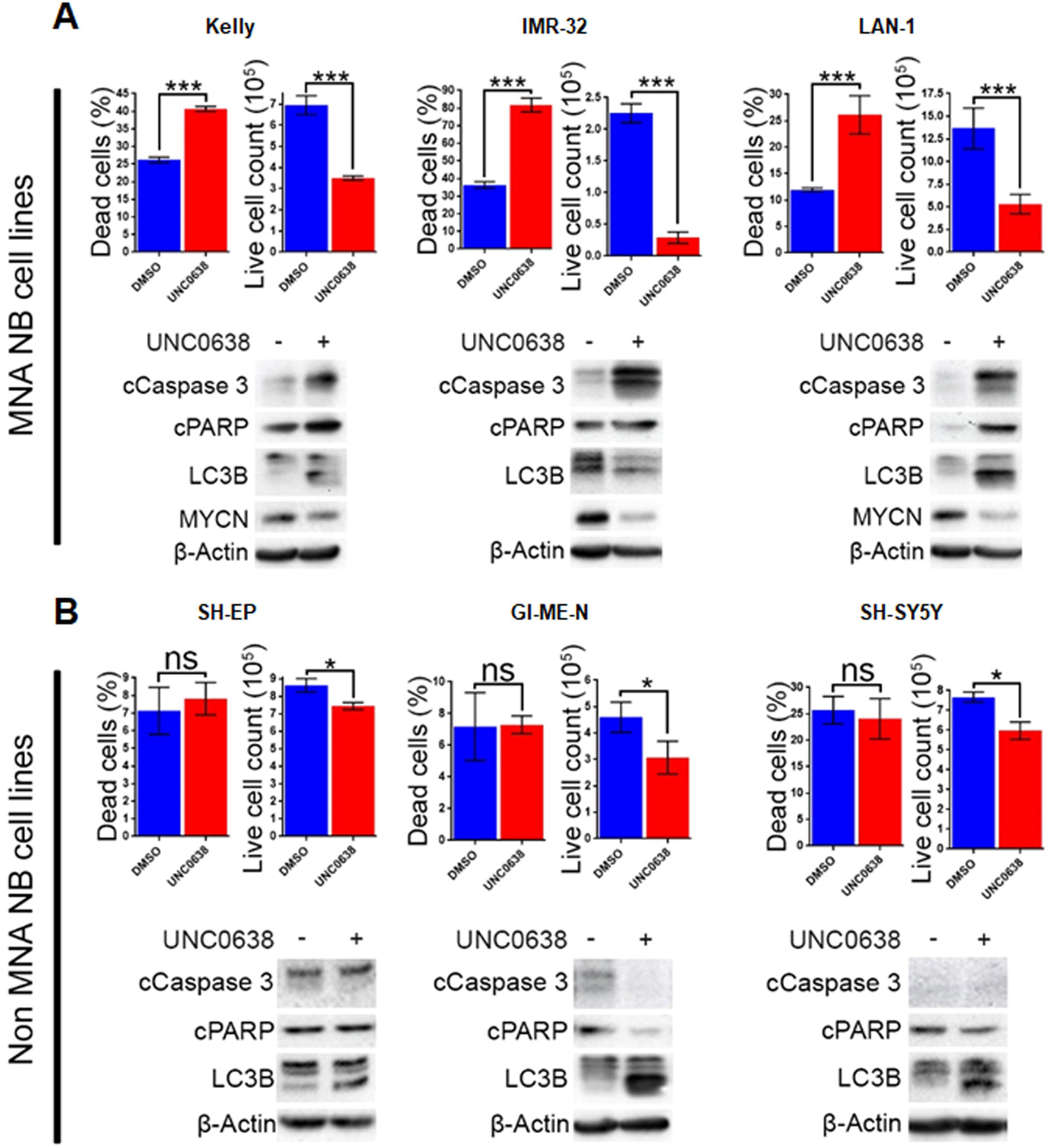
UNC0638 specifically induces apoptosis in MNA neuroblastoma cell-lines. **(A)** Floating and adherent cells from MNA cell lines were harvested and counted by trypan blue inclusion assay following 5-10 μM UNC0638 treatment for 72 hours. The left-hand graph of each cell line shows the percentage dead cells, while the righthand graph shows the live cell counts, which was used as a proxy for cell growth. Significant changes are indicated by asterisks (*** p<0.01, n=3). Error bars show the SEM. Western blots for each cell line show markers for cell death, autophagy markers and MYCN. β-Actin is used as loading control. (representative of n=3). **(B)** Live and dead cell counts and Western blots of non-MNA cells after treatment with 10μM UNC0638 for 72 hours. (* p<0.05, ns not significant, n=3)

Conversely, in the three non-MNA lines, SH-EP, GI-M-EN, and SH-SY5Y, there was no significant change in the percentage dead cells. However, there was a significant decrease in the number of live cells counted following the treatments (Figure 6B). There was no change in the apoptotic markers after UNC0638 treatment; however, there was an increase in the autophagy marker LC3B in all three non-MNA cell-lines. Together these experiments mirror the G9a depletion experiments with respect to increased apoptosis and efficacy in MNA cell-lines.

Using the S21N model, we further evaluated the requirement for MYCN over-expression in apoptosis triggered by UNC0638. Although there was no increase in the percentage of dead cells without MYCN induction, a significant 2-fold increase in dead cells was apparent in the presence of MYCN induction (Figure 7A). Immunoblotting showed that the autophagy marker LC3B was increased following UNC0638 treatment regardless of MYCN levels. The apoptosis markers, however, were only increased by UNC0638 in the presence of MYCN over-expression, further confirming the requirement for G9a activity for the survival of MYCN over-expressing cells (Figure 7B). This further supports G9a activity for targeted therapeutics in NB, especially for patients with MYCN amplification.

**Figure 7.**
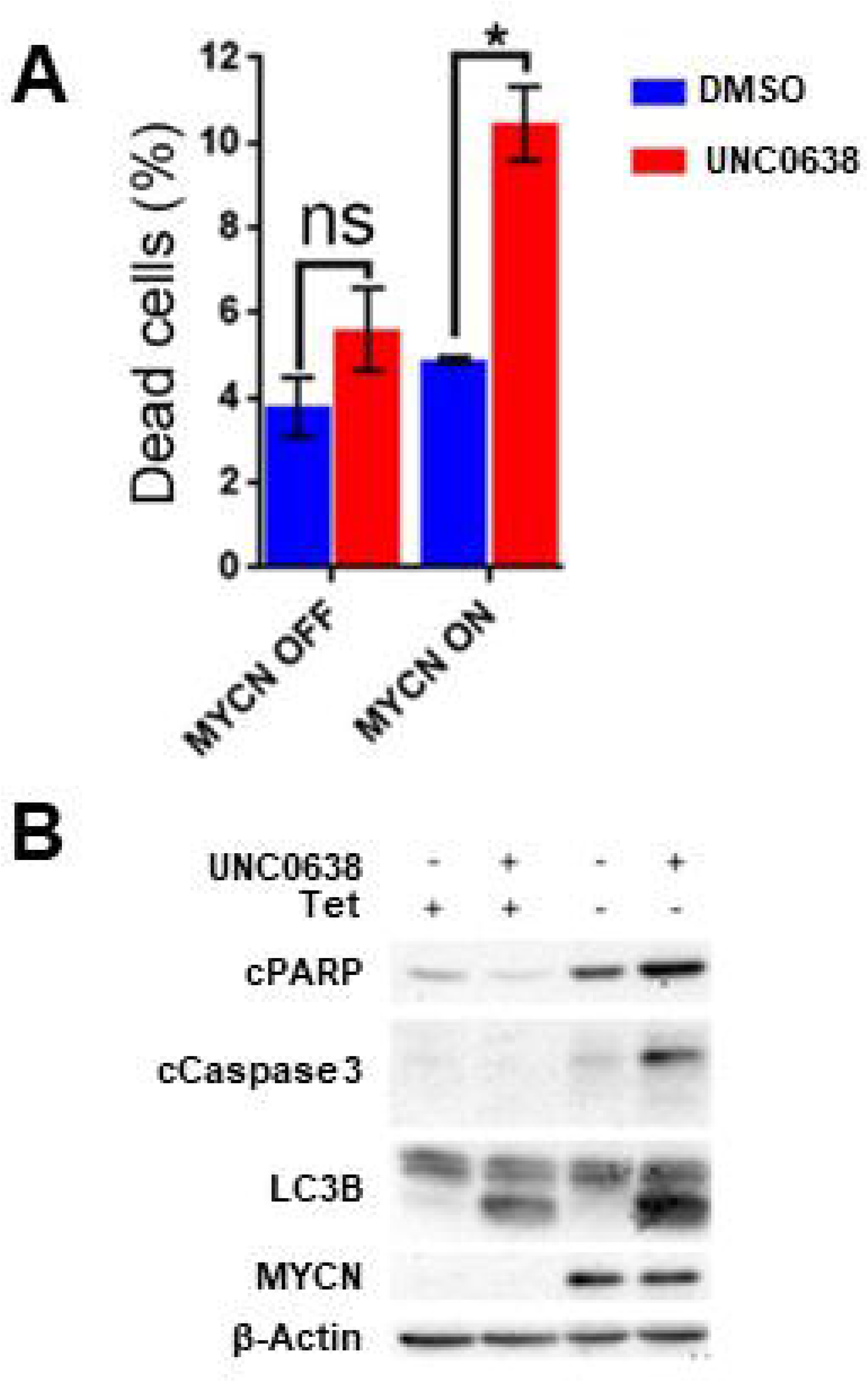
UNC0638 leads to apoptosis of MYCN overexpressing S21N cells. **(A)** Floating and adherent cells from S21N cells with and without induced MYCN were harvested and counted by trypan blue inclusion assay following 5μM UNC0638 treatment at 72 hours. The charts show the percentage dead and live cells between treated and control. Error bars are SEM. (* p<0.05, ns not significant, n=3). **(B)** Western blot of induced and uninduced S21N cells treated with 5μM UNC0638 for 72 hours.

### Identification of genes regulated by G9a inhibition in NB

We next sought insight into gene expression changes in MNA NB cells following G9a inhibition. For this, we treated LAN-1 cells with BIX-01294, as this inhibitor would facilitate comparison with previous studies on G9a in NB, including one attributing drug activity to changes in the expression of genes involved in serine metabolism. Using RNA sequencing, we identified 115 genes whose expression level was altered by more than 1.3-fold, at p<0.005. Consistent with the role of G9a in epigenetic repression, the majority of these genes were upregulated after BIX-01294 treatment, but approximately a third of affected genes were down-regulated (Figure 8A). The magnitude of gene induction was generally much higher than the changes in down-regulated genes, with 11 genes being upregulated between 10-100-fold more than vehicle treated cells. In contrast, down-regulated genes were decreased by a maximum of approximately 5-fold. We have not detected changes in the genes of the serine-glycine pathway as previously described (Ding et al. 2013) which may be due to differences in culture conditions, namely supplementation with non-essential amino acids. A full list of genes showing altered expression is given in Supplementary Table 2. Gene set enrichment analysis (GSEA) verified a profound effect on gene sets driven by the MYC family; in particular the MYCN-157 signature of genes associated with poor prognosis in NB (Valentijn et al. 2012) was profoundly affected. MYCN induced genes from this data set were downregulated by BIX-01294, whereas MYCN repressed genes were upregulated (Figure 8B). GSEA also showed that apoptosis gene sets were upregulated, together with gene sets including genes epigenetically silenced by EZH2 and histone deacetylases 1 and 3 (HDAC1 and HDAC3) (Figure 8C-D). These analyses suggest that G9a inhibition may be effective in altering oncogenic gene expression programmes in MNA NB.

**Figure 8.**
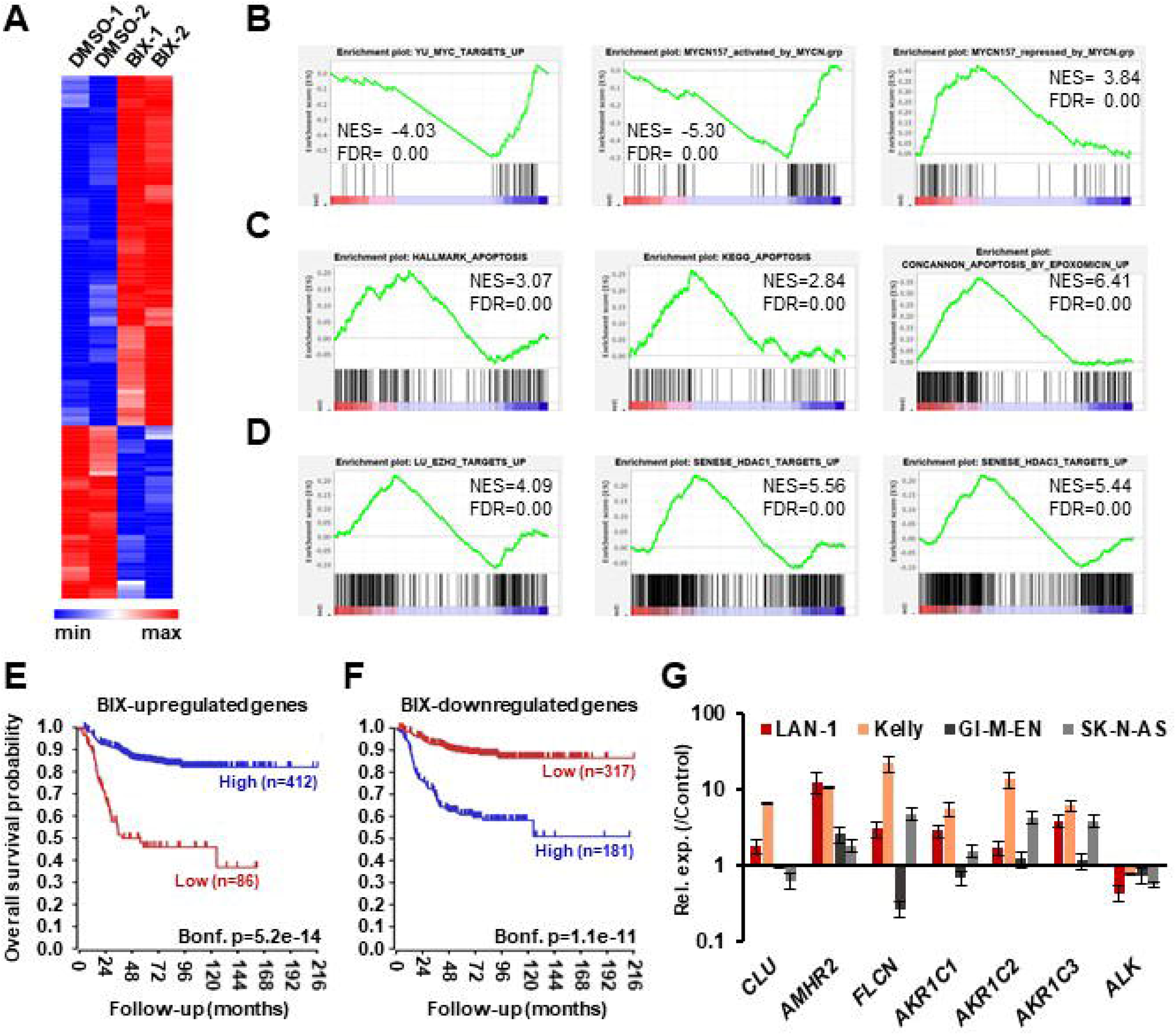
RNA sequencing of BIX-01294 treated LAN-1 cells identifies G9a regulated genes. **(A)** Heatmap of differentially expressed genes (DEGs) in biological replicate treatments of LAN-1 NB cells by 3 μM BIX-01294 for 72 hours, (p<0.005, minimum fold change 1.3). **(B)** Gene set enrichment analysis (GSEA) showing reversal of MYC/MYCN-driven transcriptomic changes. The MYCN-157 gene sets were derived from Valentijn et al. 2012. **(C)** GSEA indicating upregulation of gene sets associated with apoptosis and **(D)** repression by EZH2, HDAC1 and HDAC3. **(E)** Kaplan-Meier survival analyses showing that high expression of genes upregulated by BIX-01294 treatment in LAN-1 correlates with good prognosis in an expression data set of 498 primary NB tumours (SEQC, GSE62564), while **(F)** high expression of BIX-01294 downregulated genes correlates with poor prognosis. **(G)** Validation of DEG in MNA NB cell lines, LAN1 and Kelly, and non-MNA cells, GI-M-EN and SK-N-AS, after treatment with G9a inhibitor (5-10 μM UNC0638 for 24 hours).

In order to further assess the alteration of oncogenic potential by G9a inhibition, we constructed metagenes corresponding to BIX-01294 upregulated and downregulated genes. These metagenes act as quantifiable model genes enabling the association with prognosis of grouped up/down regulated genes. As shown in Figure 8E, high expression of BIX-01294 upregulated genes correlate with good overall outcome, and low expression of these genes correlates with poor prognosis. Conversely, high expression of BIX-01294 down-regulated genes correlates with poor overall outcome, and low expression of these genes correlates with good prognosis. This strongly suggests that G9a represses tumour suppressor genes in NB that can be reactivated by BIX-01294 treatment. The data also suggests that G9a supports activation of genes associated with NB development, including MYCN regulated genes.

Lastly, we validated some of the top genes identified after BIX-01294 treatment of LAN-1 cells in two MNA (LAN-1 and Kelly) and two non-MNA NB (GI-M-EN and SK-N-AS) cell-lines treated with UNC0638. As shown in Figure 8F, *CLU, AMHR2, FLCN*, and *AKR1C1-3* were all upregulated to varying extents following UNC0638 treatment, confirming good concordance between the actions of BIX-01294 and UNC0638. We also confirmed down-regulation of the *ALK* oncogene, revealed by our RNA sequencing, with UNC0638. The association between expression of individual genes and clinical prognosis is summarized in Supplementary Table 2, strongly suggesting that our BIX-01294/UNC0638 validated genes, *AMHR2, FLCN*, and *AKR1C1-3* are amongst many novel NB tumour suppressor genes found by our analysis. *CLU*, encoding clusterin, is already a known NB tumour suppressor gene (Chayka et al. 2009) known to be repressed by MYCN and EZH2 (Valentijn et al. 2012; Wang et al. 2012).

Taken together, our data strongly support small molecule inhibition of G9a as a therapeutic intervention for NB, in particular the MYCN-amplified subgroup.

## Discussion

Whilst the potential of targeting the epigenetic machinery for cancer therapy is increasingly recognised (Pfister and Ashworth 2017), and over-expression of G9a has been reported in many cancers (Casciello et al. 2015), targeted inhibition of G9a is relatively understudied. In the context of neuroblastoma, three papers have examined the impact of G9a knockdown and the small molecule inhibitor BIX-01294 on cell proliferation and cell death (Ke et al. 2014; Lu et al. 2013; Ding et al. 2013). However, the effect and potential benefits of targeting G9a in NB remain unclear due to limited analysis of the G9a protein in NB, activities and pathways regulated by G9a, and the absence of assessment of second-generation inhibitors of G9a towards NB cells. In this study, we have examined G9a expression in relation to NB disease stratifying factors, and also assessed three G9a inhibitors. Based on our findings, as discussed below, we propose that pharmaceutical inhibition of G9a is a viable therapeutic approach, especially for NB driven by MYCN amplification.

Our immunohistochemical analysis of G9a in NB shows for the first time that nuclear G9a is markedly increased in poorly differentiated and undifferentiated NB, as also suggested by our mRNA database mining and immunoblotting of thirteen NB cell-lines. Two other potentially critical associations were revealed by these studies, (i) elevated G9a in MNA cell-lines, and (ii) correlated expression between EZH2 and G9a in primary tumours. Interestingly, G9a has been shown to stabilise c-Myc in immune cells and thereby contribute to the regulation of inflammation (Liu et al. 2014). More recently, this association was also reported in breast cancer cell lines, with the Myc-G9a complex being crucial for Myc-mediated gene repression (Tu et al. 2018). Thus, it is possible that G9a may regulate MYCN in an analogous manner by modulating MYCNs transcriptional activity. G9a may also act similarly to PRMT5, which we have shown to directly methylate MYCN and regulate its stability at the protein level (Park et al. 2015). The need for further in-depth analyses to explore the possible interplay of G9a and MYCN proteins is underlined by our expression data, as well as functional and transcriptomic data (see below).

The correlation of G9a and EZH2 expression is also important when considering possible epigenetic therapeutics for NB. Although G9a is mainly known to catalyse H3K9 dimethylation, it can also methylate H3K27 (Tachibana et al. 2001) and it has been shown to be a key regulator of Polycomb repressor Complex 2 (Mozzetta et al. 2014). In addition, it was shown in breast cancer cells that effective gene re-expression necessitated the inhibition of both G9a and EZH2. For example, dual depletion of G9a and EZH2 dramatically increased *SPINK1* mRNA when individual depletion had no effect. This dual inhibition was also shown to increase growth inhibition over only G9a or EZH2 single inhibition (Curry et al. 2015). Given the increasing evidence for EZH2 involvement in NB, especially in tandem with MYCN (Li et al. 2018; Chen et al. 2018; Corvetta et al. 2013; Wang et al. 2012), there is clearly a rationale for deploying G9a and EZH2 inhibitors together for the treatment of NB. This is further emphasized by the reactivation of the *CLU* gene by all the G9a inhibitors shown in our studies, as it is a NB tumour suppressor gene known to be regulated by EZH2 and MYCN (Chayka et al. 2009; Valentijn et al. 2012; Wang et al. 2012).

Our evaluation of UNC0638 and UNC0642 showed that both inhibitors have a more pronounced growth-inhibitory effect on MNA NB cell-lines. Furthermore, functional analysis of genetic and pharmaceutical inhibition of G9a revealed a striking correlation between MYCN over-expression and apoptosis triggered by G9a inhibition. This clarifies to some extent the previous contradictions regarding autophagy and apoptosis resulting from G9a inhibition in NB (Ke et al. 2014; Lu et al. 2013; Ding et al. 2013). More importantly, it alludes to a synthetic lethal relationship (Kaelin Jr 2005) between G9a and MYCN expression in MNA NB. Whilst the mechanisms underlying this remain to be fully elucidated, our RNA sequencing suggests that a simultaneous combination of effects on MYCN control of gene expression and epigenetic derepression are strongly involved. G9a is also known to be involved in the DNA damage response, with UNC0638 potentiating the cytotoxicity of DNA damaging agents (Agarwal and Jackson 2016). It is therefore possible that cells over-expressing MYCN have greater replicative stress and are therefore more susceptible to an impairment/inhibition of the DNA damage response. Although G9a is also known to regulate the p53 protein by post-translational methylation (Huang et al. 2010), our data showing that the *TP53* wild-type cell line IMR-32 shows comparable sensitivity to other MNA cell-lines (containing *TP53* mutations) suggests that the p53 pathway is not directly involved in growth inhibition induced by the G9a inhibitors.

Our RNA sequencing following BIX-01294 treatment revealed upregulation of the established NB tumour suppressors *CLU*, but also other putative tumour suppressors not previously associated with NB. One example of this is the *FLCN* gene, encoding folliculin. Folliculin has been shown to regulate AMP-activated kinase (AMPK), which enables regulation of cancer cell metabolism and also autophagy (Possik et al. 2014; Yan et al. 2014). Notably, our upregulated genes also included *FNIP1* and *FNIP2*, encoding folliculin-interacting proteins 1 and 2, emphasising the potential importance of this pathway in NB tumour suppression. Other putative tumour suppressor genes include *AMHR2*, encoding Anti-Mullerian Hormone Receptor Type 2, also known as Mullerian Inhibiting Substance Type II Receptor. *AMHR2* has been shown to suppress tumorigenicity in the testes (Tanwar et al. 2012). The aldo keto-reductase 1 family genes (*AKR1C1-3*) encode steroidogenic genes which, although expressed at high levels in some cancers (Zeng et al. 2017), are also downregulated in others such as breast and gastric cancers (Frycz et al. 2016; Wenners et al. 2016). Of the down-regulated genes, several were histone genes, probably reflecting the decreased G1 to S-phase progression in cells treated with inhibitors. The *ALK* gene was also decreased, possibly as a result of decreased MYCN; ALK and MYCN are known to mutually regulate each other in NB (Hasan et al. 2013; Schonherr et al. 2012). Whilst our study does not establish a direct causal link between G9a, MYCN and ALK, it is interesting to note that the two cell-lines most sensitive to UNC0638 are representative of “ultra-high risk” NB, having both *MYCN* amplification and activating mutations of *ALK*.

In summary, this paper highlights a previously unrecognised therapeutic vulnerability of neuroblastomas with MYCN amplification to small molecule inhibitors of G9a, As MYCN is also a known driver of several other cancers, this work underlines the need for future work on these cancers with current inhibitors, and the development of next generation G9a inhibitors.

---

**Supplementary Figure 1S. Apoptosis and cell cycle analysis of Kelly cells following UNC0638 treatment. (A)** Floating and adherent cells from Kelly cells G9a were harvested and counted by trypan blue inclusion assay following G9a depletion and treatment with pan caspase inhibitor QVD 10μM for 72 hours. Error bars are SEM. (*** p<0.01, n=3). **(B)** Immunoblot showing that G9a depletion associated apoptosis is rescuable following QVD treatment. **(C)** Cell cycle analysis of Kelly cells treated with 3μM UNC0638 for 72 hours. There is a significant increase in sub G1 population, an increase in G1 arrest, and a significant reduction of cells in S phase (* p<0.05, *** p<0.01, n=3).

## 2 Conflict of Interest

The authors declare that the research was conducted in the absence of any commercial or financial relationships that could be construed as a potential conflict of interest.

## 3 Author Contributions

JB, MS, AD and ZM carried out all experiments with assistance from MK. DC provided critical reagents. KM designed the study and wrote the manuscript, with assistance from JB & MS.

## 4 Funding

This work was supported by Cancer Research UK (A12743/A21046) (to K.M.), together with the Childrens Cancer and Leukaemia Group (CCLG), Children with Cancer UK, and the Showering Fund (NHS).

## 5 Acknowledgments

We wish to thank the Genomics and Flow Cytometry facilities at the University of Bristol for technical help.

## 1 Data Availability Statement

The datasets generated for this study can be found at http://www.ebi.ac.uk/ena/data/view/PRJEB35417.

## Notes

http://www.ebi.ac.uk/ena/data/view/PRJEB35417

## REFERENCES

Agarwal, P., and S. P. Jackson. 2016. ‘G9a inhibition potentiates the anti-tumour activity of DNA double-strand break inducing agents by impairing DNA repair independent of p53 status’, Cancer Lett, 380: 467–75.

Brodeur, G. M. 2003. ‘Neuroblastoma: biological insights into a clinical enigma’, Nat Rev Cancer, 3: 203–16.

Brodeur, G. M., R. C. Seeger, M. Schwab, H. E. Varmus, and J. M. Bishop. 1984. ‘Amplification of N-myc in untreated human neuroblastomas correlates with advanced disease stage’, Science, 224: 1121.

Casciello, F., K. Windloch, F. Gannon, and J. S. Lee. 2015. ‘Functional Role of G9a Histone Methyltransferase in Cancer’, Front Immunol, 6: 487.

Chayka, O., D. Corvetta, M. Dews, A. E. Caccamo, I. Piotrowska, G. Santilli, S. Gibson, N. J. Sebire, N. Himoudi, M. D. Hogarty, J. Anderson, S. Bettuzzi, A. Thomas-Tikhonenko, and A. Sala. 2009. ‘Clusterin, a haploinsufficient tumor suppressor gene in neuroblastomas’, J Natl Cancer Inst, 101: 663–77.

Chen, L., G. Alexe, N. V. Dharia, L. Ross, A. B. Iniguez, A. S. Conway, E. J. Wang, V. Veschi, N. Lam, J. Qi, W. C. Gustafson, N. Nasholm, F. Vazquez, B. A. Weir, G. S. Cowley, L. D. Ali, S. Pantel, G. Jiang, W. F. Harrington, Y. Lee, A. Goodale, R. Lubonja, J. M. Krill-Burger, R. M. Meyers, A. Tsherniak, D. E. Root, J. E. Bradner, T. R. Golub, C. W. Roberts, W. C. Hahn, W. A. Weiss, C. J. Thiele, and K. Stegmaier. 2018. ‘CRISPR-Cas9 screen reveals a MYCN-amplified neuroblastoma dependency on EZH2’, J Clin Invest, 128: 446–62.

Cho, Hyun-Soo, John D. Kelly, Shinya Hayami, Gouji Toyokawa, Masahi Takawa, Masanori Yoshimatsu, Tatsuhiko Tsunoda, Helen I. Field, David E. Neal, Bruce A. J. Ponder, Yusuke Nakamura, and Ryuji Hamamoto. 2011. ‘Enhanced Expression of EHMT2 Is Involved in the Proliferation of Cancer Cells through Negative Regulation of SIAH1’, Neoplasia, 13: 676–IN10.

Corvetta, D., O. Chayka, S. Gherardi, C. W. D’Acunto, S. Cantilena, E. Valli, I. Piotrowska, G. Perini, and A. Sala. 2013. ‘Physical interaction between MYCN oncogene and polycomb repressive complex 2 (PRC2) in neuroblastoma: functional and therapeutic implications’, J Biol Chem, 288: 8332–41.

Curry, E., I. Green, N. Chapman-Rothe, E. Shamsaei, S. Kandil, F. L. Cherblanc, L. Payne, E. Bell, T. Ganesh, N. Srimongkolpithak, J. Caron, F. Li, A. G. Uren, J. P. Snyder, M. Vedadi, M. J. Fuchter, and R. Brown. 2015. ‘Dual EZH2 and EHMT2 histone methyltransferase inhibition increases biological efficacy in breast cancer cells’, Clin Epigenetics, 7: 84.

Ding, J., T. Li, X. Wang, E. Zhao, J. H. Choi, L. Yang, Y. Zha, Z. Dong, S. Huang, J. M. Asara, H. Cui, and H. F. Ding. 2013. ‘The histone H3 methyltransferase G9A epigenetically activates the serine-glycine synthesis pathway to sustain cancer cell survival and proliferation’, Cell Metab, 18: 896–907.

Frycz, B. A., D. Murawa, M. Borejsza-Wysocki, M. Wichtowski, A. Spychala, R. Marciniak, P. Murawa, M. Drews, and P. P. Jagodzinski. 2016. ‘Transcript level of AKR1C3 is down-regulated in gastric cancer’, Biochem Cell Biol, 94: 138–46.

Gherardi, S., E. Valli, D. Erriquez, and G. Perini. 2013. ‘MYCN-mediated transcriptional repression in neuroblastoma: the other side of the coin’, Front Oncol, 3: 42.

Hasan, M. K., A. Nafady, A. Takatori, S. Kishida, M. Ohira, Y. Suenaga, S. Hossain, J. Akter, A. Ogura, Y. Nakamura, K. Kadomatsu, and A. Nakagawara. 2013. ‘ALK is a MYCN target gene and regulates cell migration and invasion in neuroblastoma’, Sci Rep, 3: 3450.

Helin, K., and D. Dhanak. 2013. ‘Chromatin proteins and modifications as drug targets’, Nature, 502: 480–8.

Huang, Jing, Jean Dorsey, Sergei Chuikov, Xinyue Zhang, Thomas Jenuwein, Danny Reinberg, and Shelley L. Berger. 2010. ‘G9a and Glp Methylate Lysine 373 in the Tumor Suppressor p53’, The Journal of Biological Chemistry, 285: 9636–41.

Iraci, N., D. Diolaiti, A. Papa, A. Porro, E. Valli, S. Gherardi, S. Herold, M. Eilers, R. Bernardoni, G. Della Valle, and G. Perini. 2011. ‘A SP1/MIZ1/MYCN repression complex recruits HDAC1 at the TRKA and p75NTR promoters and affects neuroblastoma malignancy by inhibiting the cell response to NGF’, Cancer Res, 71: 404–12.

Kaelin Jr, William G. 2005. ‘The Concept of Synthetic Lethality in the Context of Anticancer Therapy’, Nature Reviews Cancer, 5: 689.

Ke, X. X., D. Zhang, S. Zhu, Q. Xia, Z. Xiang, and H. Cui. 2014. ‘Inhibition of H3K9 methyltransferase G9a repressed cell proliferation and induced autophagy in neuroblastoma cells’, PLoS One, 9: e106962.

Ke, X. X. 2019. ‘Correction: Inhibition of H3K9 Methyltransferase G9a Repressed Cell Proliferation and Induced Autophagy in Neuroblastoma Cells’, PLoS One, 14: e0213135.

Kondo, Yutaka, Lanlan Shen, Seiji Suzuki, Tsuyoshi Kurokawa, Kazuo Masuko, Yasuhito Tanaka, Hideaki Kato, Yoshiki Mizuno, Masamichi Yokoe, Fuminaka Sugauchi, Noboru Hirashima, Etsuro Orito, Hirotaka Osada, Ryuzo Ueda, Yi Guo, Xinli Chen, J. Issa Jean-Pierre, and Yoshitaka Sekido. 2007. ‘Alterations of DNA methylation and histone modifications contribute to gene silencing in hepatocellular carcinomas’, Hepatology Research, 37: 974–83.

Kubicek, S., R. J. O’Sullivan, E. M. August, E. R. Hickey, Q. Zhang, M. L. Teodoro, S. Rea, K. Mechtler, J. A. Kowalski, C. A. Homon, T. A. Kelly, and T. Jenuwein. 2007. ‘Reversal of H3K9me2 by a small-molecule inhibitor for the G9a histone methyltransferase’, Mol Cell, 25: 473–81.

Lee, David Y., Jeffrey P. Northrop, Min-Hao Kuo, and Michael R. Stallcup. 2006. ‘HISTONE H3 LYSINE 9 METHYLTRANSFERASE G9a IS A TRANSCRIPTIONAL COACTIVATOR FOR NUCLEAR RECEPTORS’, The Journal of Biological Chemistry, 281: 8476–85.

Lee, Jason S., Yunho Kim, Jinhyuk Bhin, Hi-Jai R. Shin, Hye Jin Nam, Seung Hoon Lee, Jong-Bok Yoon, Olivier Binda, Or Gozani, Daehee Hwang, and Sung Hee Baek. 2011. ‘Hypoxia-induced methylation of a pontin chromatin remodeling factor’, Proceedings of the National Academy of Sciences, 108: 13510.

Lee, Jason S., Yunho Kim, Ik Soo Kim, Bogyou Kim, Hee June Choi, Ji Min Lee, Hi-Jai R. Shin, Jung Hwa Kim, Ji-Young Kim, Sang-Beom Seo, Ho Lee, Olivier Binda, Or Gozani, Gregg L. Semenza, Minhyung Kim, Keun Il Kim, Daehee Hwang, and Sung Hee Baek. 2010. ‘Negative Regulation of Hypoxic Responses via Induced Reptin Methylation’, Molecular Cell, 39: 71–85.

Li, Zhenghao, Hisanori Takenobu, Amallia Nuggetsiana Setyawati, Nobuhiro Akita, Masayuki Haruta, Shunpei Satoh, Yoshitaka Shinno, Koji Chikaraishi, Kyosuke Mukae, Jesmin Akter, Ryuichi P. Sugino, Atsuko Nakazawa, Akira Nakagawara, Hiroyuki Aburatani, Miki Ohira, and Takehiko Kamijo. 2018. ‘EZH2 regulates neuroblastoma cell differentiation via NTRK1 promoter epigenetic modifications’, Oncogene, 37: 2714–27.

Liu, C., Y. Yu, F. Liu, X. Wei, J. A. Wrobel, H. P. Gunawardena, L. Zhou, J. Jin, and X. Chen. 2014. ‘A chromatin activity-based chemoproteomic approach reveals a transcriptional repressome for gene-specific silencing’, Nat Commun, 5: 5733.

Liu, F., D. Barsyte-Lovejoy, F. Li, Y. Xiong, V. Korboukh, X. P. Huang, A. Allali-Hassani, W. P. Janzen, B. L. Roth, S. V. Frye, C. H. Arrowsmith, P. J. Brown, M. Vedadi, and J. Jin. 2013. ‘Discovery of an in vivo chemical probe of the lysine methyltransferases G9a and GLP’, J Med Chem, 56: 8931–42.

Lu, Z., Y. Tian, H. R. Salwen, A. Chlenski, L. A. Godley, J. U. Raj, and Q. Yang. 2013. ‘Histonelysine methyltransferase EHMT2 is involved in proliferation, apoptosis, cell invasion, and DNA methylation of human neuroblastoma cells’, Anticancer Drugs, 24: 484–93.

Maris, J. M., M. D. Hogarty, R. Bagatell, and S. L. Cohn. 2007. ‘Neuroblastoma’, Lancet, 369: 2106–20.

Mossë, Yalë P., Marci Laudenslager, Luca Longo, Kristina A. Cole, Andrew Wood, Edward F. Attiyeh, Michael J. Laquaglia, Rachel Sennett, Jill E. Lynch, Patrizia Perri, Geneviève Laureys, Frank Speleman, Hakon Hakonarson, Ali Torkamani, Nicholas J. Schork, Garrett M. Brodeur, Gian Paolo Tonini, Eric Rappaport, Marcella Devoto, and John M. Maris. 2008. ‘Identification of ALK as the Major Familial Neuroblastoma Predisposition Gene’, Nature, 455: 930–35.

Mozzetta, C., J. Pontis, L. Fritsch, P. Robin, M. Portoso, C. Proux, R. Margueron, and S. Ait-Si-Ali. 2014. ‘The histone H3 lysine 9 methyltransferases G9a and GLP regulate polycomb repressive complex 2-mediated gene silencing’, Mol Cell, 53: 277–89.

Park, Ji Hyun, Marianna Szemes, Gabriella Cunha Vieira, Zsombor Melegh, Sally Malik, Kate J. Heesom, Laura Von Wallwitz-Freitas, Alexander Greenhough, Keith W. Brown, Y. George Zheng, Daniel Catchpoole, Michael J. Deery, and Karim Malik. 2015. ‘Protein arginine methyltransferase 5 is a key regulator of the MYCN oncoprotein in neuroblastoma cells’, Molecular Oncology, 9: 617–27.

Pfister, S. X., and A. Ashworth. 2017. ‘Marked for death: targeting epigenetic changes in cancer’, Nat Rev Drug Discov, 16: 241–63.

Possik, E., Z. Jalali, Y. Nouet, M. Yan, M. C. Gingras, K. Schmeisser, L. Panaite, F. Dupuy, D. Kharitidi, L. Chotard, R. G. Jones, D. H. Hall, and A. Pause. 2014. ‘Folliculin regulates ampk-dependent autophagy and metabolic stress survival’, PLoS Genet, 10: e1004273.

Schonherr, C., K. Ruuth, S. Kamaraj, C. L. Wang, H. L. Yang, V. Combaret, A. Djos, T. Martinsson, J. G. Christensen, R. H. Palmer, and B. Hallberg. 2012. ‘Anaplastic Lymphoma Kinase (ALK) regulates initiation of transcription of MYCN in neuroblastoma cells’, Oncogene, 31: 5193–200.

Shinkai, Y., and M. Tachibana. 2011. ‘H3K9 methyltransferase G9a and the related molecule GLP’, Genes Dev, 25: 781–8.

Su, Zhenqiang, Hong Fang, Huixiao Hong, Leming Shi, Wenqian Zhang, Wenwei Zhang, Yanyan Zhang, Zirui Dong, Lee J. Lancashire, Marina Bessarabova, Xi Yang, Baitang Ning, Binsheng Gong, Joe Meehan, Joshua Xu, Weigong Ge, Roger Perkins, Matthias Fischer, and Weida Tong. 2014. ‘An investigation of biomarkers derived from legacy microarray data for their utility in the RNA-seq era’, Genome Biology, 15: 3273.

Szemes, M., A. Greenhough, Z. Melegh, S. Malik, A. Yuksel, D. Catchpoole, K. Gallacher, M. Kollareddy, J. H. Park, and K. Malik. 2018. ‘Wnt Signalling Drives Context-Dependent Differentiation or Proliferation in Neuroblastoma’, Neoplasia, 20: 335–50.

Tachibana, M., K. Sugimoto, T. Fukushima, and Y. Shinkai. 2001. ‘Set domain-containing protein, G9a, is a novel lysine-preferring mammalian histone methyltransferase with hyperactivity and specific selectivity to lysines 9 and 27 of histone H3’, J Biol Chem, 276: 25309–17.

Tachibana, Makoto, Kenji Sugimoto, Masami Nozaki, Jun Ueda, Tsutomu Ohta, Misao Ohki, Mikiko Fukuda, Naoki Takeda, Hiroyuki Niida, Hiroyuki Kato, and Yoichi Shinkai. 2002. ‘G9a histone methyltransferase plays a dominant role in euchromatic histone H3 lysine 9 methylation and is essential for early embryogenesis’, Genes & Development, 16: 1779–91.

Tanwar, P. S., A. E. Commandeur, L. Zhang, M. M. Taketo, and J. M. Teixeira. 2012. ‘The Mullerian inhibiting substance type 2 receptor suppresses tumorigenesis in testes with sustained beta-catenin signaling’, Carcinogenesis, 33: 2351–61.

Tu, W. B., Y. J. Shiah, C. Lourenco, P. J. Mullen, D. Dingar, C. Redel, A. Tamachi, W. Ba-Alawi, A. Aman, R. Al-Awar, D. W. Cescon, B. Haibe-Kains, C. H. Arrowsmith, B. Raught, P. C. Boutros, and L. Z. Penn. 2018. ‘MYC Interacts with the G9a Histone Methyltransferase to Drive Transcriptional Repression and Tumorigenesis’, Cancer Cell, 34: 579–95 e8.

Valentijn, L. J., J. Koster, F. Haneveld, R. A. Aissa, P. van Sluis, M. E. Broekmans, J. J. Molenaar, J. van Nes, and R. Versteeg. 2012. ‘Functional MYCN signature predicts outcome of neuroblastoma irrespective of MYCN amplification’, Proc Natl Acad Sci U S A, 109: 19190–5.

Vedadi, M., D. Barsyte-Lovejoy, F. Liu, S. Rival-Gervier, A. Allali-Hassani, V. Labrie, T. J. Wigle, P. A. Dimaggio, G. A. Wasney, A. Siarheyeva, A. Dong, W. Tempel, S. C. Wang, X. Chen, I. Chau, T. J. Mangano, X. P. Huang, C. D. Simpson, S. G. Pattenden, J. L. Norris, D. B. Kireev, A. Tripathy, A. Edwards, B. L. Roth, W. P. Janzen, B. A. Garcia, A. Petronis, J. Ellis, P. J. Brown, S. V. Frye, C. H. Arrowsmith, and J. Jin. 2011. ‘A chemical probe selectively inhibits G9a and GLP methyltransferase activity in cells’, Nat Chem Biol, 7: 566–74.

Wang, C., Z. Liu, C. W. Woo, Z. Li, L. Wang, J. S. Wei, V. E. Marquez, S. E. Bates, Q. Jin, J. Khan, K. Ge, and C. J. Thiele. 2012. ‘EZH2 Mediates epigenetic silencing of neuroblastoma suppressor genes CASZ1, CLU, RUNX3, and NGFR’, Cancer Res, 72: 315–24.

Wen, Bo, Hao Wu, Yoichi Shinkai, Rafael A. Irizarry, and Andrew P. Feinberg. 2009. ‘Large histone H3 lysine 9 dimethylated chromatin blocks distinguish differentiated from embryonic stem cells’, Nature Genetics, 41: 246.

Wenners, A., F. Hartmann, A. Jochens, A. M. Roemer, I. Alkatout, W. Klapper, M. van Mackelenbergh, C. Mundhenke, W. Jonat, and M. Bauer. 2016. ‘Stromal markers AKR1C1 and AKR1C2 are prognostic factors in primary human breast cancer’, Int J Clin Oncol, 21: 548–56.

Westermark, U. K., M. Wilhelm, A. Frenzel, and M. A. Henriksson. 2011. ‘The MYCN oncogene and differentiation in neuroblastoma’, Seminars in Cancer Biology, 21: 256–66.

Yan, M., M. C. Gingras, E. A. Dunlop, Y. Nouet, F. Dupuy, Z. Jalali, E. Possik, B. J. Coull, D. Kharitidi, A. B. Dydensborg, B. Faubert, M. Kamps, S. Sabourin, R. S. Preston, D. M. Davies, T. Roughead, L. Chotard, M. A. van Steensel, R. Jones, A. R. Tee, and A. Pause. 2014. ‘The tumor suppressor folliculin regulates AMPK-dependent metabolic transformation’, J Clin Invest, 124: 2640–50.

Zeng, C. M., L. L. Chang, M. D. Ying, J. Cao, Q. J. He, H. Zhu, and B. Yang. 2017. ‘Aldo-Keto Reductase AKR1C1-AKR1C 4: Functions, Regulation, and Intervention for Anti-cancer Therapy’, Front Pharmacol, 8: 119.

Zhang, Jie, Pengxing He, Yong Xi, Meiyu Geng, Yi Chen, and Jian Ding. 2014. ‘Down-regulation of G9a triggers DNA damage response and inhibits colorectal cancer cells proliferation’, Oncotarget, 6(5): 2917–2927.

