## Supplementary figures and images for "Increased efficacy of histone methyltransferase G9a inhibitors against MYCN-amplified Neuroblastoma"

### Supplementary Figure 1.

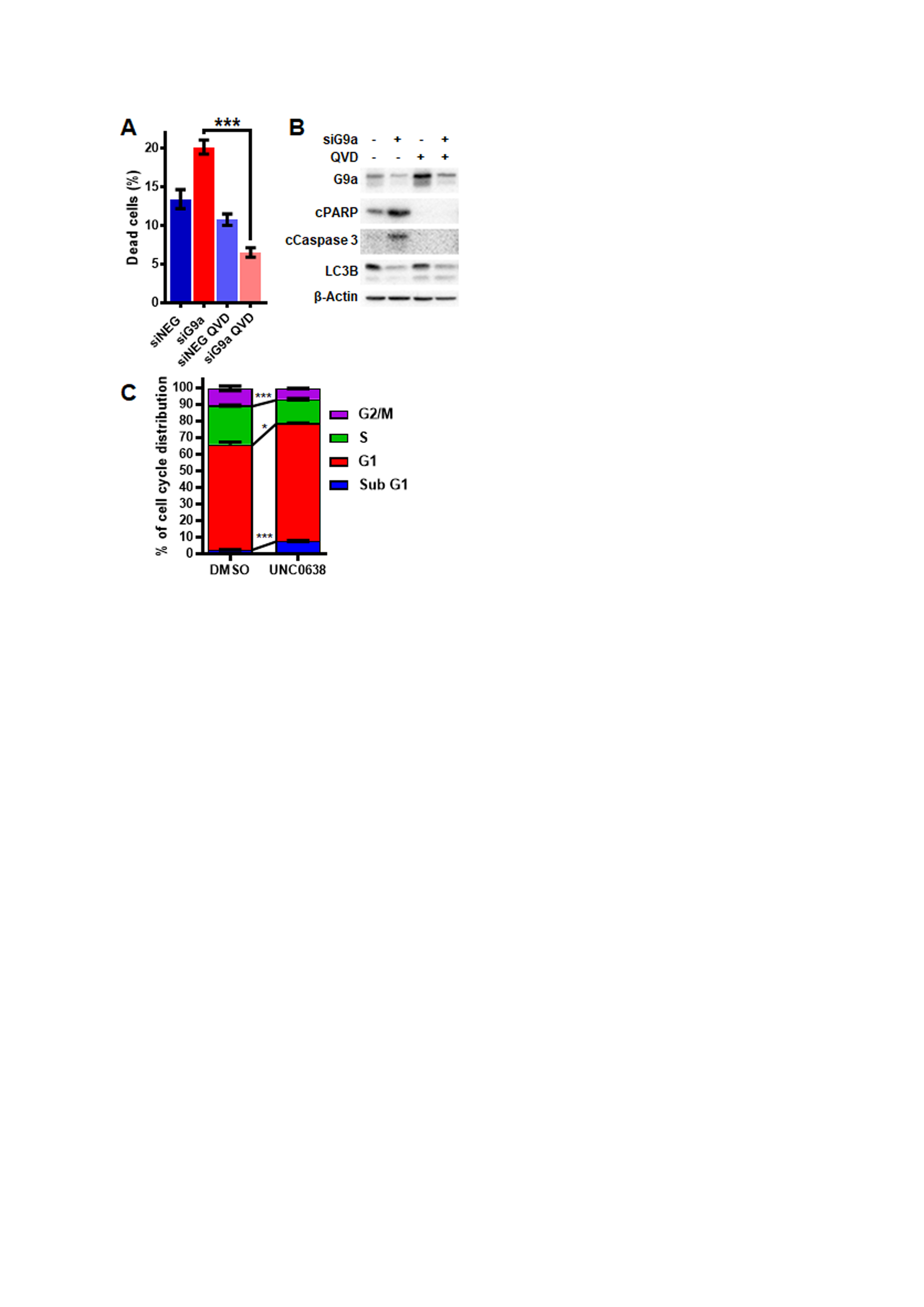
