## Supplementary table 1. for "Increased efficacy of histone methyltransferase G9a inhibitors against MYCN-amplified Neuroblastoma"

| Gene | Fold Change | FDR | Kaplan-Meier analysis (SEQC) | |
|---|---|---|---|---|
| | | | High Expression | Bonferroni p |
| *AMHR2* | 100.30 | 2.23E-06 | good prognosis | 1.75E-05 |
| *BHLHE40* | 39.96 | 2.61E-05 | good prognosis | 3.51E-08 |
| *KLHDC8A* | 38.36 | 4.27E-04 | good prognosis | 4.78E-13 |
| *PMEL* | 32.75 | 4.29E-11 | | ns |
| *ACP5* | 30.86 | 1.41E-05 | good prognosis | 5.62E-08 |
| *AKR1C3* | 26.30 | 2.80E-21 | good prognosis | 7.32E-07 |
| *DIRAS2* | 20.55 | 1.75E-11 | good prognosis | 2.46E-12 |
| *AKR1C2* | 15.03 | 8.21E-15 | good prognosis | 1.01E-19 |
| *CTD-2516F10.2* | 12.51 | 7.60E-05 | good prognosis | 3.90E-07 |
| *RENBP* | 12.24 | 1.81E-03 | good prognosis | 9.99E-08 |
| *PLP1* | 9.59 | 9.46E-09 | good prognosis | 9.67E-08 |
| *UAP1L1* | 7.38 | 5.24E-17 | | ns |
| *AKR1C1* | 6.81 | 1.16E-03 | good prognosis | 2.71E-24 |
| *PRKG2* | 6.76 | 2.56E-03 | | ns |
| *HOPX* | 5.96 | 3.40E-09 | good prognosis | 1.47E-05 |
| *CYB561* | 5.62 | 1.56E-05 | good prognosis | 4.04E-15 |
| *COLEC11* | 5.41 | 1.66E-13 | poor prognosis | 3.00E-05 |
| *FTL* | 4.89 | 5.71E-32 | poor prognosis | 2.60E-01 |
| *HMOX1* | 4.80 | 2.13E-03 | | ns |
| *BRI3* | 4.47 | 2.75E-08 | poor prognosis | 2.95E-03 |
| *DNM3* | 4.31 | 2.78E-05 | good prognosis | 8.49E-13 |
| *GCHFR* | 3.78 | 5.32E-11 | poor prognosis | 5.73E-08 |
| *SC5D* | 3.75 | 1.09E-11 | | ns |
| *ALDOC* | 3.73 | 7.60E-05 | good prognosis | 2.92E-17 |
| *SLC7A14* | 3.71 | 1.87E-04 | | ns |
| *HMGCS1* | 3.45 | 4.65E-09 | | ns |
| *MSMO1* | 3.23 | 4.91E-11 | poor prognosis | 1.25E-09 |
| *INSIG1* | 3.05 | 1.28E-13 | | ns |
| *CLU* | 2.95 | 1.96E-07 | good prognosis | 7.03E-14 |
| *ARMCX3* | 2.85 | 3.54E-07 | good prognosis | 1.58E-06 |
| *SQSTM1* | 2.84 | 2.54E-12 | good prognosis | 6.44E-11 |
| *FLCN* | 2.79 | 2.78E-05 | good prognosis | 9.69E-22 |
| *FDFT1* | 2.78 | 9.95E-14 | poor prognosis | 5.87E-11 |
| *STMN2* | 2.77 | 4.11E-07 | good prognosis | 5.19E-15 |
| *PPM1H* | 2.72 | 8.45E-05 | poor prognosis | 3.77E-02 |
| *ARMCX1* | 2.67 | 1.05E-04 | good prognosis | 2.90E-02 |
| *GPIHBP1* | 2.66 | 1.99E-05 | good prognosis | 3.10E-07 |
| *DDIT3* | 2.66 | 4.38E-03 | good prognosis | 1.97E-11 |
| *SLC30A1* | 2.56 | 3.06E-05 | | ns |
| *LONRF2* | 2.47 | 3.69E-03 | good prognosis | 3.14E-23 |
| *PIM3* | 2.41 | 5.06E-04 | poor prognosis | 4.75E-08 |
| *CD63* | 2.35 | 7.67E-06 | | ns |
| *ZBTB38* | 2.29 | 2.91E-04 | good prognosis | 6.49E-16 |
| *AHNAK* | 2.26 | 2.72E-08 | good prognosis | 2.23E-04 |
| *SQLE* | 2.25 | 1.22E-05 | poor prognosis | 5.12E-17 |
| *GDE1* | 2.22 | 3.06E-10 | poor prognosis | 5.45E-05 |
| *ALCAM* | 2.21 | 1.49E-03 | good prognosis | 5.12E-20 |
| *NCR3LG1* | 2.20 | 1.87E-04 | poor prognosis | 5.32E-12 |

| | | | | |
|---|---|---|---|---|
| *ASAH1* | 2.20 | 3.24E-04 | good prognosis | 2.98E-10 |
| *FNIP1* | 2.19 | 9.00E-04 | | ns |
| *SHC1* | 2.19 | 6.44E-07 | good prognosis | 1.32E-03 |
| *SFRP1* | 2.17 | 1.14E-08 | good prognosis | 1.79E-14 |
| *DUSP3* | 2.17 | 3.31E-05 | good prognosis | 4.42E-10 |
| *SH3BP5* | 2.11 | 4.94E-05 | good prognosis | 1.73E-13 |
| *KLHL24* | 2.10 | 1.37E-07 | good prognosis | 9.45E-03 |
| *GNPDA1* | 2.08 | 8.16E-06 | poor prognosis | 2.70E-17 |
| *FNIP2* | 2.06 | 9.22E-04 | good prognosis | 4.83E-06 |
| *SLC31A1* | 2.05 | 1.27E-05 | good prognosis | 4.01E-03 |
| *GSTM3* | 2.03 | 3.62E-03 | good prognosis | 1.23E-06 |
| *HMGCR* | 2.01 | 1.92E-06 | poor prognosis | 1.04E-05 |
| *KLHDC8B* | 1.99 | 1.16E-03 | good prognosis | 1.50E-02 |
| *IDI1* | 1.96 | 2.89E-04 | poor prognosis | 1.99E-03 |
| *MIER1* | 1.93 | 3.61E-03 | good prognosis | 1.20E-14 |
| *KIAA0930* | 1.92 | 1.80E-03 | good prognosis | 3.83E-20 |
| *ACAT2* | 1.91 | 6.46E-04 | poor prognosis | 7.28E-04 |
| *DHCR24* | 1.90 | 4.43E-04 | | ns |
| *RB1CC1* | 1.87 | 2.05E-06 | | ns |
| *SLC4A8* | 1.84 | 6.51E-05 | good prognosis | 5.49E-04 |
| *GOLGA4* | 1.83 | 6.88E-05 | poor prognosis | 1.33E-05 |
| *QPRT* | 1.79 | 6.63E-04 | poor prognosis | 7.78E-13 |
| *STARD4* | 1.76 | 2.81E-05 | good prognosis | 3.20E-02 |
| *PSAP* | 1.76 | 1.61E-03 | good prognosis | 1.27E-14 |
| *DSE* | 1.75 | 3.42E-03 | good prognosis | 1.75E-04 |
| *SCG2* | 1.74 | 2.13E-03 | good prognosis | 3.86E-18 |
| *GPAM* | 1.72 | 1.87E-03 | poor prognosis | 1.47E-05 |
| *LRRD1* | 1.72 | 1.68E-03 | good prognosis | 1.14E-15 |
| *AFF4* | 1.70 | 2.75E-03 | | ns |
| *FLRT3* | 0.59 | 3.07E-06 | poor prognosis | 2.93E-05 |
| *DPYSL5* | 0.58 | 4.07E-03 | | ns |
| *PHOX2A* | 0.57 | 2.89E-04 | | ns |
| *PTGFRN* | 0.57 | 2.26E-03 | | ns |
| *HIST1H2BJ* | 0.56 | 3.50E-14 | poor prognosis | 6.00E-06 |
| *H2AFX* | 0.56 | 4.38E-03 | poor prognosis | 5.41E-20 |
| *NNAT* | 0.55 | 2.68E-03 | good prognosis | 3.27E-08 |
| *MYBL2* | 0.55 | 6.89E-04 | poor prognosis | 1.27E-14 |
| *CADM1* | 0.55 | 1.07E-04 | good prognosis | 3.67E-15 |
| *TEX15* | 0.55 | 1.09E-04 | poor prognosis | 3.60E-24 |
| *HIST1H2AL* | 0.54 | 3.10E-11 | | ns |
| *FAM64A* | 0.54 | 1.12E-03 | poor prognosis | 8.10E-17 |
| *HIST1H2AB* | 0.54 | 6.61E-07 | poor prognosis | 2.21E-04 |
| *AURKB* | 0.54 | 3.63E-03 | poor prognosis | 4.74E-16 |
| *HIST1H2BN* | 0.53 | 8.58E-04 | poor prognosis | 7.05E-03 |
| *FAM83D* | 0.53 | 1.58E-03 | poor prognosis | 3.81E-30 |
| *HIST1H4I* | 0.52 | 7.18E-13 | poor prognosis | 9.29E-10 |
| *NCAPD2* | 0.52 | 7.42E-06 | poor prognosis | 1.12E-17 |
| *RGS4* | 0.51 | 2.16E-06 | good prognosis | 3.72E-05 |
| *VCAN* | 0.51 | 1.68E-06 | poor prognosis | 8.99E-03 |
| *HIST1H2AG* | 0.48 | 3.09E-22 | poor prognosis | 4.33E-07 |

| | | | | |
|---|---|---|---|---|
| *ZNF850* | 0.44 | 2.57E-04 | poor prognosis | 5.70E-30 |
| *COL4A2* | 0.41 | 6.29E-07 | good prognosis | 2.67E-02 |
| *FN1* | 0.40 | 5.16E-10 | | ns |
| *CCDC144NL-AS1* | 0.37 | 3.41E-03 | na | na |
| *CTD-2314G24.2* | 0.36 | 1.89E-03 | na | na |
| *TENM4* | 0.35 | 4.94E-05 | poor prognosis | 1.28E-14 |
| *FBN2* | 0.34 | 2.57E-04 | | ns |
| *DLK1* | 0.33 | 8.42E-08 | poor prognosis | 3.68E-08 |
| *ARHGAP36* | 0.32 | 2.04E-20 | good prognosis | 1.43E-09 |
| *FAT3* | 0.32 | 3.66E-06 | good prognosis | 3.72E-04 |
| *SHISA2* | 0.31 | 9.65E-05 | poor prognosis | 5.30E-11 |
| *FIGF* | 0.29 | 3.27E-18 | poor prognosis | 1.51E-13 |
| *GFRA1* | 0.27 | 3.47E-03 | good prognosis | 1.52E-03 |
| *ALK* | 0.24 | 6.93E-04 | poor prognosis | 3.29E-07 |
| *NPNT* | 0.22 | 1.80E-03 | good prognosis | 2.78E-07 |
| *CDH11* | 0.20 | 6.63E-04 | good prognosis | 1.02E-05 |
| *IGSF1* | 0.19 | 4.03E-03 | good prognosis | 9.37E-06 |
